# Decay in transcriptional information flow is a hallmark of cellular aging

**DOI:** 10.1101/2025.10.30.685689

**Authors:** Brooke Emison, Christopher W. Lynn, Andrew Mugler, Fabrisia Ambrosio, Purushottam Dixit

## Abstract

Aging is marked by the progressive loss of cellular function, yet the organizing principles underlying this decline remain unclear. Although molecular fingerprints of aging are diverse, many converge on disruption of the interrelated and overlapping communication networks that coordinate molecular activity. Here, we apply information theory to quantify age-related corruption in gene regulation by modeling regulatory interactions between transcription factors (TFs) and their target genes (TGs) as a multi-input multi-output communication channel. Using an analytically tractable probabilistic model and single-cell RNA-sequencing data from multiple tissues, we find that the mutual information (a measure of information transfer) between TFs and TGs declines with age across all ten tissues analyzed, establishing loss of regulatory information transmission as a hallmark of aging. Structural analysis of the regulatory network reveals that aging degrades communication primarily through input distribution mismatch, reflecting a loss of coordinated TF activity, rather than channel corruption, or the inability of TFs to reliably activate or inhibit their targets. This mismatch is caused by increased network centralization and loss of stabilizing feedback motifs, leading to reduced robustness to random perturbations. Notably, *in silico* upregulation of a small set of TFs restores youthful information transfer and gene expression levels, suggesting that targeted reinforcement of key regulatory nodes may rejuvenate aged networks.

## Introduction

Aging is the single greatest risk factor for multiple chronic diseases [1]. Despite its diverse manifestations, a unifying feature is the progressive loss of cellular function. Notably, many molecular fingerprints of cellular aging [2-4] (e.g., increase in oxidative stress, protein damage, genomic instability, and epigenetic drift) converge on altered intracellular communication through changes in gene regulatory networks (GRNs).

GRNs are essential in maintaining cellular homeostasis via their role in response to intracellular stress and environmental cues. Prior work on bulk transcriptomics in *C. elegans*, mice, and humans revealed that stress-related genes increase while metabolic and growth genes decrease with age [5-8]. Further analysis of bulk transcriptomics across multiple tissues across human subjects showed that predictive gene/gene interactions weakened with age, especially between genes belonging to different functional modules, implying a loss of coordination among biological programs [9, 10]. However, correlation structures inferred from bulk data may reflect subject heterogeneity in regulatory programs rather than true age-associated decoupling.

Single-cell studies have extended these analyses to study how gene expression noise changes with age. Because most genes are expressed at only a few copies per cell, random fluctuations preclude fine-tuned regulation. Pioneering work by Bahar *et al*. showed increased transcriptional noise in aged cardiomyocytes [11]. Later work extended these analyses to different animals and tissues [12-18]. Our previous work in muscle cells showed that aging reduced correlation between transcription factors and their targets, with the extent of decoupling heterogeneously distributed across hallmarks of aging [19]. Yet, evidence remains mixed as to whether age-related transcript and protein noise increases are gene- and tissue-dependent [20]. An improved understanding of how aging alters the architecture of GRNs across the organism could reveal unifying principles of cellular decline and help identify new therapeutic targets to restore system-wide coordination.

Since precise regulation of gene expression requires accurate communication between transcription factors (TFs) and their target genes (TGs), information theory provides a natural framework to quantify communication fidelity in GRNs [21, 22]. Unlike previous correlation-based approaches that only capture linear and pairwise interactions, information theoretic quantification can account for nonlinear and multi-input–multi-output dependencies, making information theory ideally suited to assess age-related information loss in GRNs. Indeed, information theory has been used to quantify the ability of signaling [23] and gene regulatory networks [22] to discern extracellular ligand concentrations and spatiotemporal variation. Notably, it has been shown how cells can use multidimensional and temporally varying outputs [24] as well as cellular heterogeneity to maximize information about their environments [25]. In the context of aging, recent work [26] has hypothesized that cellular aging can be viewed through the lens of loss of communication in gene regulation and blurring of cellular identity.

However, there has been no rigorous quantification of potential aging-related deterioration in gene regulatory networks. This is in part due to the difficulty in estimating information theoretic quantities such as mutual information (MI) and entropy from high-dimensional single-cell data, as direct quantification from data scale poorly with data dimensionality [27]. Analytically tractable probabilistic models overcome this limitation by providing an interpretable representation of the underlying data distribution. These models allow closed-form expressions for MI, enabling quantitative comparisons of information flow across tissues and ages. Here, we develop such a model, inspired by the statistical physics of spin glasses [28, 29], to compute MI between TFs and their TGs. Using a multi-tissue public single-cell atlas [30], we show that MI systematically decreases with age, revealing loss of information transmission as a hallmark of aging.

Our framework distinguishes two modes of information decay [31]: corruption of the regulatory channel (i.e., loss of the ability of TFs to reliably activate or repress their targets) versus mismatch of TF input distributions (i.e., altered TF expression patterns that compromise the inputs received by targets). We find that input mismatch predominates as the cause of aging-related deterioration in gene regulation, which is also reflected in large scale structural reorganization of the TF/TF interaction networks, including reduced feedback motifs, broader out-degree distributions, and diminished stability to random perturbations. From a biological perspective, this means that aging disrupts the coordinated balance of TF activities, rather than the fundamental ability of these factors to regulate their targets. These findings imply that, similar to Yamanaka factors [32] that restore pluripotency in differentiated cells, a more youthful network phenotype can be achieved by rebalancing aged TF activities [33-35]. Indeed, we identified a subset of TFs whose upregulation restored youthful communication patterns, suggesting potential rejuvenation strategies. Together, these findings position loss of information transmission within GRNs as a universal and potentially reversible hallmark of aging.

## Results

### Non-equilibrium spin-glass model accurately captures gene expression profiles

To quantify how TFs control gene expression, we developed a probabilistic model of the GRN (Figure 1). Briefly, the transcriptome was divided into TFs (taken from a database [36]) and their TGs. TFs regulate other genes, including other TFs, while TGs are defined as genes that are regulated but do not regulate others. Both TFs and TGs are treated as binary variables (active = 1, inactive = 0). The probabilities of gene activities and TF activities were expressed as:

**Figure 1.**
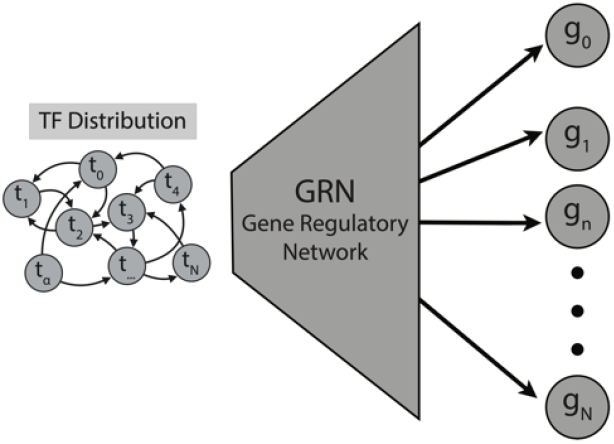
Schematic of the computational model of the gene regulatory network. A recurrent network of interactions (left, TF Distribution) between transcription factors generates the transcription factor (TF) activity distribution (left). A feedforward network represents the gene regulatory network (GRN) that predicts gene activities from TF activities.

and

In Eq. 1, is the magnitude of influence of TF on TG, captures the constitutive activity of the TG, and represents the binary vector of activities of all TFs. Similarly, in Eq. 2, is the influence of TF on TF and represents activities of all TFs except . While the probability of TG activation was modeled as a single-layer feedforward network (Eq. 1), TF activities were governed by a recurrent interaction network that generates a non-equilibrium stationary distribution over simultaneous activities of all TFs. Together, these two distributions define a joint probabilistic generative model for TF and TG activity.

We trained this model using single-cell transcriptomic data from the Tabula Muris Senis atlas [30] across 10 different tissues from young (3 months) and aged (24 months) mice, focusing on SmartSeq data to minimize dropout noise [37]. Model parameters were inferred using Lasso regularization to prevent overfitting and accurately reproduce mean activities and pairwise covariations (see SI section 1, SI Tables 1 and 2). The model reproduced both mean expression and correlation structures with high fidelity (Pearson r = 0.97 for TFs, 0.98 for TGs; 0.89 for TF– TF, 0.87 for TG–TG, and 0.76 for TF–TG correlations for limb muscles; Figure 2, see SI Figures 1-10). Therefore, this minimal formulation enabled realistic *in silico* transcriptomes and allowed exact probability computations of gene activities within the GRN (Eq. 1).

**Figure 2.**
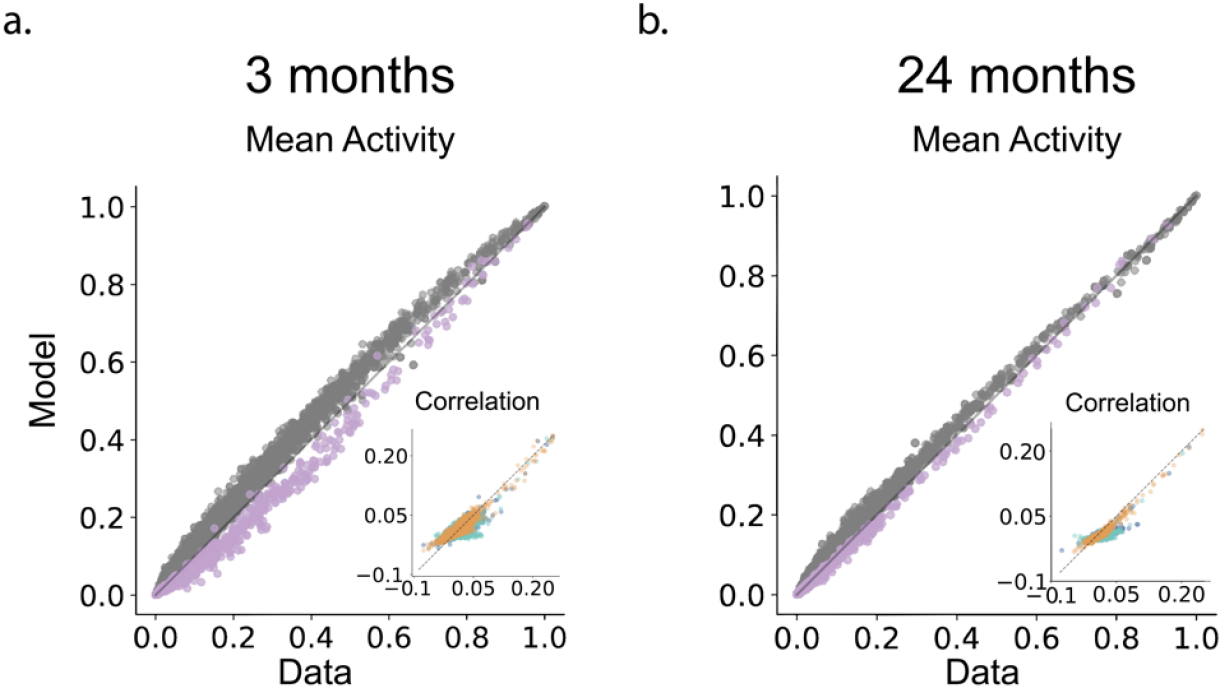
Computational model captures gene expression statistics. **a**. A comparison between mean activities of TGs (gray) and TFs (purple) as observed in the data (x-axis) and as predicted by the model (y-axis) for limb muscle data collected on 3-month-old mice. **b**. Same as a. for data collected on muscle cells from 24-month-old mice. Insets: A comparison between TG/TG correlation (blue), TF/TF correlation (orange) and TF/TG correlation (green) as observed in the data (x-axis) and as predicted by the model (y-axis). The dashed black line represents the x = y line.

### Communication fidelity in gene regulatory networks decreases across all tissues

Communication fidelity in an input/output network can be rigorously quantified using mutual information (MI). The MI *I*_*i*_, between a target gene “*i*” (the output) and all transcription factors (the input) is defined as [31]

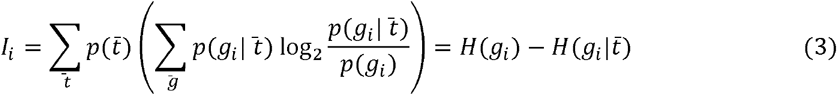

In Eq. 3,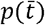is the stationary distribution over TFs, obtained by simulating Eq. 2 as a Markov chain 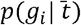 is the probability of activation of a gene as a function of input TFs given by Eq. 1 and *p*(*g*_*i*_)is the marginalized probability that gene is active, averaging over the activity of all transcription factors.

The MI has several salient features. First, the MI is agnostic to the nature of relationship between inputs and outputs (linear or non-linear etc.) and only depends on the extent of conditional dependence. Second, the MI is always non-negative and treats activating and inhibitory influences on an equal footing; zero MI indicates that the output is statistically independent of the input: 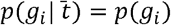. Third, MI naturally accounts for how inputs regulate the output (encoded by the conditional distribution 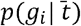 and whether the input distribution (encoded by 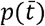 ) aligns with the input/output relationship. Finally, the MI also offers an intuitive interpretation of information transduction in terms of reduction in uncertainty. The MI can be expressed as the reduction in the entropy of the output *H*(*g*_*i*_) due to the knowledge of the input 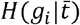. If knowing the input offers no new insight about the output, we have 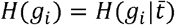 and *I*_*i*_ = 0. In contrast, if knowing the input allows us to fully determine the output, we have 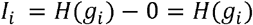.Therefore, MI is bounded by the entropy of the output itself. Specifically, the MI is maximum when given no information about TFs, we know nothing about the gene activity and we reach near certainty once the transcription factor activities are specified. In the binarized form, the maximum value of gene-specific mutual information is 1 bit.

We used Eq. 3 to numerically compute the MI between every gene and all transcription factors using age- and tissue-specific models learned from single cell RNAseq data (see SI Section 2 for details, SI Table 3, SI Figures 11-20). Notably, as shown for the example of limb muscle cells, the mutual information varied significantly between genes, ranging from close to 0 (TFs do not predict TG activity) to very close to 1 bit (knowledge of TFs predicts TG activity to near certainty). This suggests that the ability of TFs to control their target genes is heterogeneously distributed; some genes were tightly regulated by TFs while others may be constitutively expressed regardless of TF activities.

Notably, MI was on average lower in muscle cells from mice 24 months of age compared to muscle cells from mice at 3 months of age (Figure 3a, gray circles, SI Section 2). This was true not only at the level of the entire gene expression program (average over all genes), but also at the level of individual genes. This suggests that aging led to a systematic loss of regulatory control across all genes. Figure 3b shows that the fraction of genes with a decrease in mutual information was higher than 50% for all tissues considered in the study, highlighting decay in information transfer in the GRN is a hallmark of cellular aging.

**Figure 3.**
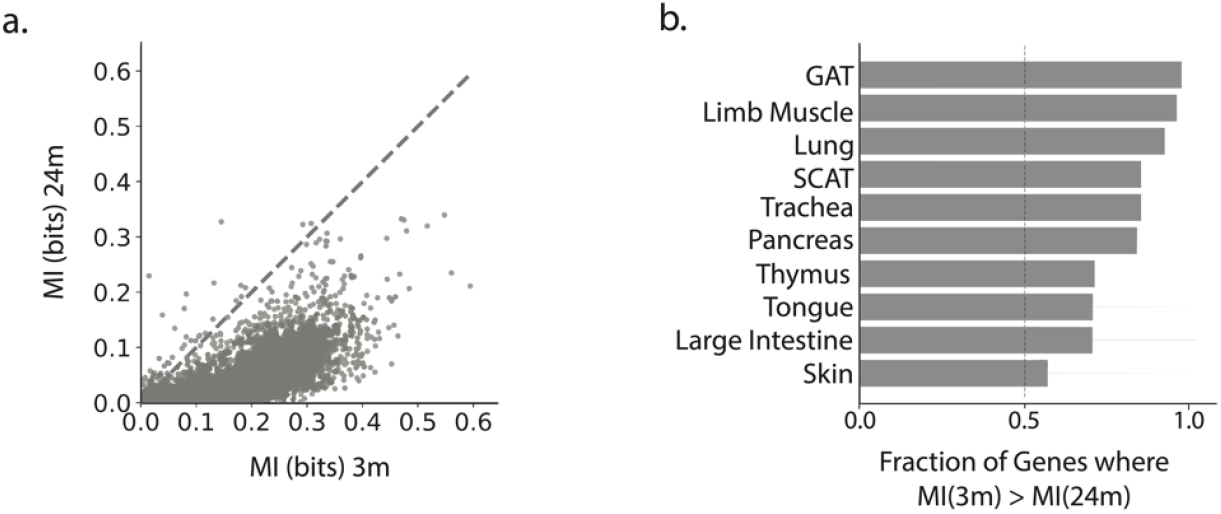
Comparison of mutual information in GRNs between ages. **a**. Comparison of mutual informatio between genes and transcription factors (each circle represents one gene) in mouse limb muscle cells collected from 3-month-old mice (x-axis) and 24-month-old mice (y-axis). **b**. The fraction of all genes where mutual information between gene activity and transcription factors was greater in cells harvested from 3-month-old mice compared to cells from 24-month-old mice across all tissues considered. Abbreviations: MI= mutual information; GAT= gonadal adipose tissue; SCAT= subcutaneous adipose tissue.

### Input mismatch, not channel corruption causes decay in information transduction

Two principal mechanisms can reduce MI (Figure 4a) [31]: (i) Channel corruption: The GRN itself becomes less responsive and/or TF binding has reduced influence on gene activation and (ii) Input distribution mismatch: TF activity distributions shift away from the sensitive regime of the regulatory input–output relationship, even if the underlying GRN remains intact. To evaluate the relative contribution of these two mechanisms, we performed heterochronic *in silico* simulations by exchanging components between young (3-month) and aged (24-month) models (Figure 4b, SI section 2, SI Table, 2, and SI Figures 11-20). Swapping the aged GRN (Eq. 1) with the young one modestly increased MI (from 0.020 to 0.036 bits per gene, ∼36% of the youthful value), while swapping the aged input distribution produced a much larger increase (to 0.062 bits per gene, ∼63 % of the youthful value, Figure 4c). These effects varied across tissues: rejuvenating the input distribution most improved MI in muscle and adipose tissue, whereas rejuvenating the GRN had larger effects in thymus and lung (Figure 4d). Overall, however, input mismatch played a predominant role in deterioration of information flow. In 8 out of 10 tissues studied (Wilcoxon signed-rank *p* = 0.03), rejuvenating input distribution led to a higher restoration of MI (on an average 85% of the youthful value), compared to rejuvenating the GRN (on an average 64% of the youthful value). These results show that aging predominantly impairs information flow through input mismatch, rather than channel corruption.

**Figure 4.**
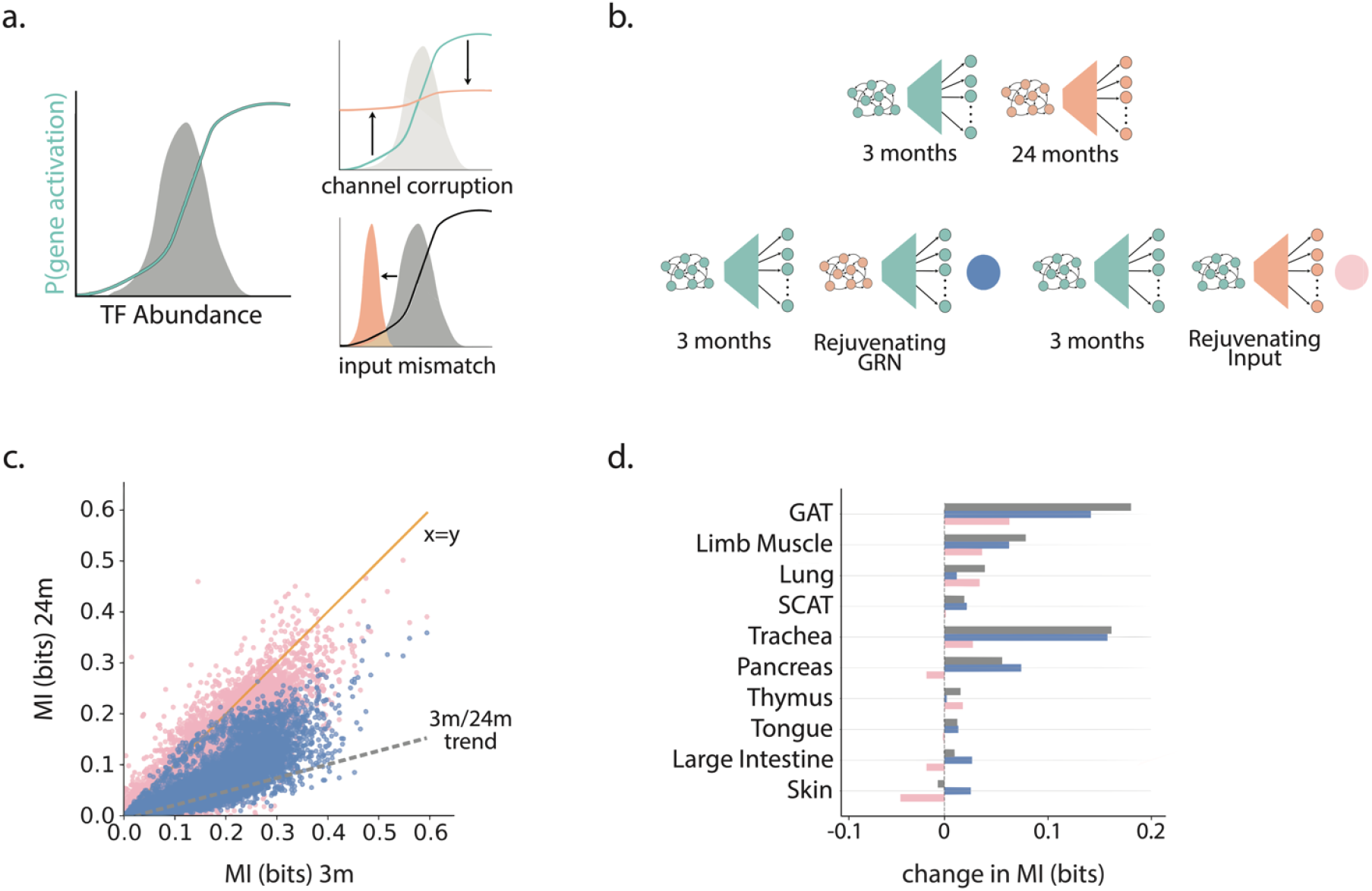
Structural causes of decay in information transfer. **a**. (left) Mutual information between input and output is high when the input distribution (TF distribution, gray) matches the sensitive region of the input/output relationship. (right): There are two modes of reducing mutual information. Channel corruption: Input distribution may not change appreciably but the input/output relationship becomes insensitive to the input. Input mismatch: The input/output channel may remain intact, but the input distribution no longer aligns with the sensitive region of the input/output relationship. **b**. A schematic showing heterochronic *in silico* simulations. Top: Mutual information (gray circles in Figure 3a) is calculated using input distribution and the gene regulatory network inferred using data from the same age (green for 3 months, orange for 24 months). Bottom left: Computational experiment comparing mutual information (blue circles) at 3 months (green and green) and in an *in silico* network comprising a 24-month input distribution (orange) and a 3-month gene regulatory network (green). Bottom right: Computational experiment comparing mutual information (pink circles) at 3 months (green and green) and in an *in silico* network comprising a 24-month gene regulatory network (orange) and a 3-month input distribution (green). **c**. Mutual information from *in silico* simulations. Blue circles represent a scenario where the gene regulatory network in 24-month-old mice was swapped with a 3-month-old network. Pink circles represent a scenario where the input distribution in 24-month-old mice was swapped with a 3-month-old input distribution. Gray dashed line represents a linear fit to the comparison between 3-month-old and 24-month-old mice (Figure 3a). **d**. The average change in mutual information between 3-and 24-month-old mice after swapping the input distribution (pink) and the gene regulatory network (blue) compared to the baseline difference (gray) across all tissues considered.

### Structural changes and destabilization of TF-TF interaction network with age

Given the central importance of the input distribution 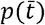 in governing information transfer in GRNs (Figure 4), we examined how the recurrent network generating the input distribution over simultaneous activities of all TFs reorganizes with age. In complex systems, aging or accumulated damage can shift control toward a few highly connected nodes [38], creating hub-dominated architectures prone to failure. Analysis of the out-degree distribution of the TF-TF interaction network revealed a marked shift toward centralization in 24-month networks compared to 3-month networks (Figure 5a); after controlling for the average density of interactions, aged networks showed a heavier tail, indicating that a smaller subset of hub TFs controlled most other TFs.

**Figure 5.**
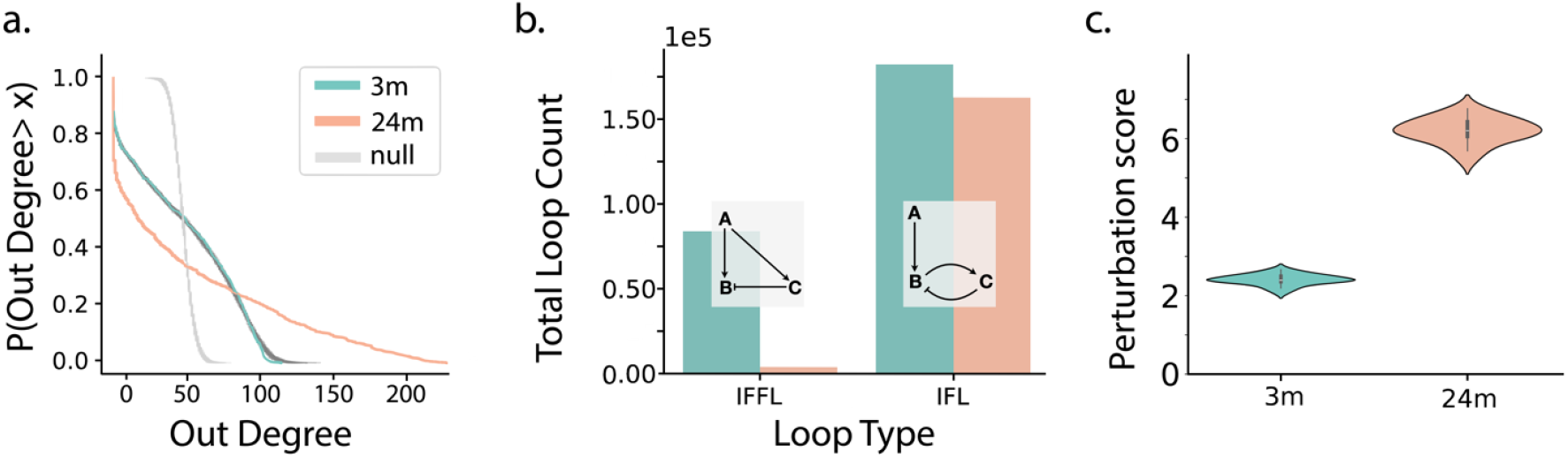
Structural properties of the inferred TF-TF interaction network. **a**. The inverse cumulative out degre distribution for transcription factors within the TF/TF interaction network constructed using cells from 3-month-ol mice (green) and 24-month-old mice (orange). The sparsity of the 3-month-old network was adjusted to match the 24-month-old network by randomly removing edges. Gray lines represent an Erdos-Renyi type null model generated by randomly assigning edges between nodes. **b**. The statistics of adaptive motifs: incoherent feedforward loops (IFFLs) and integral feedback loops (IFLs) in TF/TF networks learned from 3-month-old (green) and 24-month-old (orange) mice. **c**. The predicted change in mean TF activities (perturbation score, see SI) after randomly perturbing th transcription factor network in 3-month-old and 24-month-old mice. All analyses are shown for cells harvested from limb muscles.

We further quantified the abundance of stabilizing motifs, incoherent feedforward loops (IFFLs) and integral feedback loops (IFLs), which buffer stochastic perturbations [39]. Both IFFLs and IFLs are regulatory motifs that are employed in adaptation. When considered in isolation, in an adapting motif the downstream regulated element (“B” in Figure 5b) transiently responds to a change in the input “A” but adapts back to its baseline value even if the input is not removed. Notably, these motifs also allow the output to sense the relative (not absolute) change in the input. IFLs and IFFLs are regularly invoked to understand adaptation in signaling networks. Moreover, these loops are known to be overrepresented in bacterial gene regulatory networks as well [40]. Both motifs were significantly depleted at 24 months (10 out of 10 tissues had a depletion for both motifs, Wilcoxon signed-rank for both IFLs and IFFLs, Figure 5b). Simulations of the model confirmed that aged TF networks were more fragile: random perturbations in activities of TFs produced larger changes in TF activities at 24 months than in 3 months (Figure 5c). These findings point to loss of stabilizing structure and increased fragility of TF control networks with age, a finding that was consistent across all tissues analyzed (SI Section 3, SI Table 4, SI Figures 21-44). These results also suggest that aged tissue may undergo drift in gene expression.

Given the age-dependent centralization of the TF-TF network, we next asked whether activating a small subset of TFs could restore information flow. To identify hub TFs that control information flow, we used *in silico* TF knock-ins in the 24-month skeletal muscle network, by setting each TF’s activity to 1 (always on) one at a time, and then we computed MI using Markov Chain Monte Carlo (MCMC) sampling. Most *in silico* TF activations had no or negative effects, but a subset improved MI substantially (Figure 6a). Many of these TFs, including *Foxq1* and members of the *Ebf* family, have been implicated in muscle maintenance and regeneration (see discussion below). Simultaneously activating just the top 5 TFs: Ebf2, *Ebf1, Aebp1*, Elf3, and Foxq1, (out of nearly a 1000 TFs) increased MI from 0.020 to 0.053 bits per gene, recovering more than half the youthful information transfer (0.099 bits). This improvement exceeded the sum of individual effects (0.043 bits based on sum of individual contributions versus 0.053 bits per gene), indicating positive cooperativity. Together, these analyses reveal that aged GRNs may retain latent capacity for coordinated reactivation, and that targeted upregulation of key TFs can partially rejuvenate communication fidelity.

**Figure 6.**
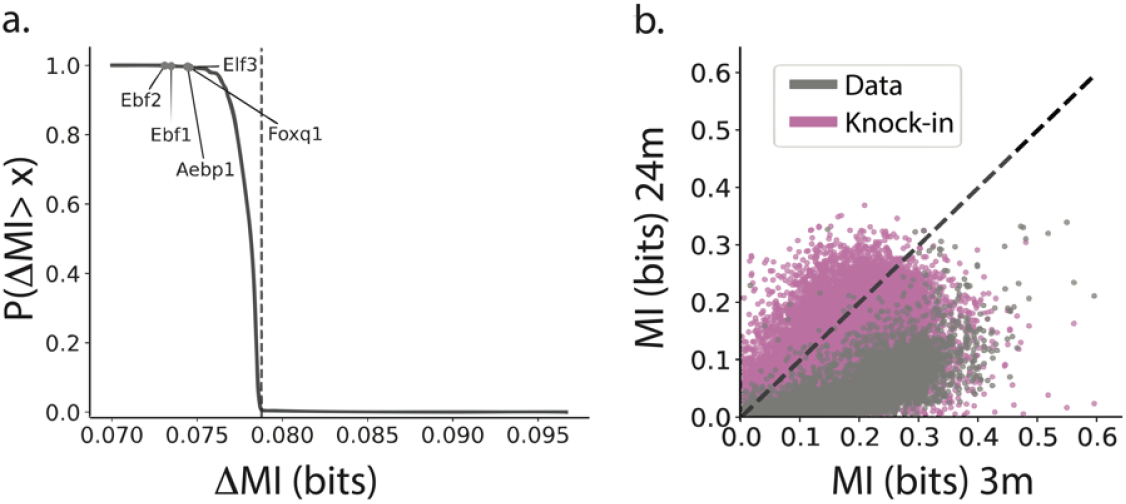
A small number of transcription factors can restore information flow. **a**. TFs are sorted according to their ability to improve the MI in gene regulatory networks at 24 months. An inverse cumulative distribution of individual TF ability to increase the MI at 24 months. Zero corresponds to MI of the 3-month network. **a**. A scatter plot of MI between genes and transcription factors (every gene is a single circle) compared between cells from 3-month-old mice (x-axis) and 24-month-old mice (y-axis). Gray circles represent original data comparison, pink circles represent a comparison when the 5 TFs, *Ebf2, Ebf1, Aebp1, Elf3*, and *Foxq1*, were knocked in *in silico* in the 24-month network.

## Discussion

In this study, we showed that cellular aging corrupts information flow across gene regulatory networks. Specifically, the mutual information between transcription factors and their target genes was consistently lower in cells isolated from old compared to young mice across all tissues analyzed in this work. This establishes loss of communication fidelity as a robust and quantitative hallmark of aging. Strikingly, in most tissues, this dysregulation arose primarily from changes in the TF input distribution, the probability distribution of joint activity of all regulatory factors, rather than from deterioration of the channel itself, i.e., the capacity of TFs to activate or repress their targets. Structural analysis of the recurrent TF networks that generate these inputs revealed age-associated centralization and loss of stability, suggesting that GRNs become hub-dominated and fragile with age.

Our framework also identified key TFs whose upregulation can re-establish youthful information flow. In limb muscles, for instance, *Foxq1* emerged as a strong candidate rejuvenation factor. Although its role in muscle aging is not fully understood, *Foxq1* is downregulated in senescent cells, and its overexpression reduces senescence while enhancing proliferation, migration, and viability of mesenchymal stem cells *in vitro* and *in vivo* [41]. Similarly, the *Ebf* family (*Ebf1/3*) acts as essential cofactors for *MyoD*, enforcing muscle-specific transcription by activating the calcium efflux pump *Serca1*. Loss of *Ebf* activity could impair Ca^2+^ homeostasis and excitation– contraction coupling, key processes in sarcopenia [42]. Together, these findings suggest that restoring communication fidelity through targeted TF modulation may represent a tractable strategy to counteract age-related muscle decline.

Beyond GRNs, our information-theoretic framework offers a general lens to study aging across multiple organizational scales. At the systems level, information theory can quantify age-related losses in cell-cell communication and tissue coordination. For example, Razban *et al*. [43] recently applied spin-glass modeling to fMRI data and found that aged human brains exhibit weaker and less integrated connectivity across regions. Similar approaches could extend to immune or endocrine communication, such as cytokine signaling during infection or regeneration. At the molecular level, our prior work showed that the IGF/FoxO pathway, a canonical longevity axis, operates near channel capacity in immortalized cells [44]. Whether such near optimal signaling is preserved or eroded in aged tissues remains an open question. More broadly, this framework may apply to pathologies that originate from dysregulated gene regulation, including cancer [45], neurodegeneration [46], and radiation injury [47], where identifying communication bottlenecks could reveal therapeutic targets.

Finally, we note several limitations. First, our analysis relies on single-cell RNA-sequencing data, which are subject to technical noise, dropout, and incomplete detection of lowly expressed TFs. The TF-target relationships we use are curated from existing databases that may omit relevant regulators or context-specific interactions. While the large number of cells enables robust detection of global differences between young and old GRNs, MI estimates for individual TF–TG pairs remain uncertain. Second, the inferred network structures are based on statistical dependencies in gene expression, which do not necessarily imply causation. Future extensions could integrate TF-DNA binding data [48], perturb-seq [49], or other multiomic assays to enhance interpretability and mechanistic resolution. Finally, our current model treats TF activity as binary; incorporating graded or time-dependent regulatory effects could capture richer dynamic behaviors, including transient adaptation or hysteresis in aging networks.

Collectively, these results reveal that aging reduces the information-processing capacity of gene regulatory networks through input mismatch, network centralization, and loss of stabilizing motifs. Yet, this decline is partially reversible, whereby the aged transcriptional landscape retains latent potential for coordinated reactivation through a small set of master regulators. By linking information theory, network structure, and rejuvenation potential, our framework provides a quantitative foundation for studying how molecular communication breaks down, and how it might be restored, with age.

## Acknowledgments

PD and BE acknowledge the support from R35GM142547. FA and AM acknowledge the support from R01AG082739. CWL acknowledges the support from R35GM160188.

## Supplemental Information

All scripts used in this manuscript can be found at

https://github.com/emisbrooke/InFlow/tree/main

## Section 1. Handling the data and model fitting

### 1.1 Data filtering

To assess information transduction in gene regulatory networks, we used the Tabula Muris Senis single cell atlas [30], which provides single cell transcriptomics data across multiple tissues and ages. The dataset includes both fluorescence-activated cell sorting (FACS, followed by smartseq2) and microfluidics droplet data. We focus our analysis on the smarseq2 data, which has a significantly lower dropout rate for genes compared to droplet-based methods [37], albeit with fewer cells. We used the processed data available at https://figshare.com/projects/Tabula_Muris_Senis/64982. The method for pre-processing is found in [30] and a jupyter notebook outlining the method in detail can be found at https://github.com/czbiohub-sf/tabula-muris-senis/tree/master. The data comprises gene counts in cells from mice that are 3m and 24m in age over 23 tissues. To exclude rarely expressed genes, we filter the dataset to include only genes that are seen in more than 3 cells in both age groups (∼ = 0.3 % ). Moreover, we only consider tissues that include at least 800 cells and non-zero mean abundances of at least 800 TFs (obtained from the humanTF dataset [36], mapped to orthologous mouse genes [50]) in both 3m and 24m. This led to 10 (out of 23) tissues used in our analysis: GAT, Limb Muscle, Lung, SCAT, Trachea, Pancreas, Thymus, Tongue, Large Intestine, and Skin.

### 1.1 Data formatting

The data is binarized such that each cell is represented by a vector of length *N*_genes_ where each element *n* is 1 if the *n*^th^ gene is present in that cell and 0 otherwise. The genes are then separated into to categories, transcription factors (TFs) and target genes (TGs). A gene is considered a TF if it is found in a large, curated compendium of TFs [36]. All other genes are considered TGs.

### 1.2 Description of the model

To model the probabilistic regulation of gene expression by transcription factors, we employ a simple biophysically motivated formulation inspired by spin-glass models of interacting binary variables. This model can be represented in two compartments, one describing how TFs interact with each other and another that describes how TFs regulate TGs. We denote TFs by Greek subscripts *α, β*, … and TGs by roman subscripts *i,j*, … We denote TFs by *t*_*α*_ = 0/1 and TGs by *g*_*i*_ = 0/1 (0 for inactive, 1 for active). We define the conditional probability of the activation of TG *i* given the activity of all TGs (denoted by a vector) 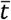 as

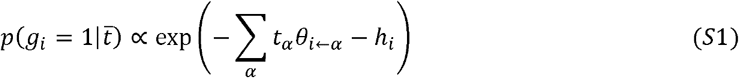

where *θ*_*i* ← *α*_ represents the effective coupling (regulatory strength) from *t*_*α*_ to *t*_*i*_ and *h*_*i*_ is a gene-specific bias term. Similarly, we define the conditional probability of the activation of *t*_*α*_ given transcription factor configuration 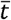 as

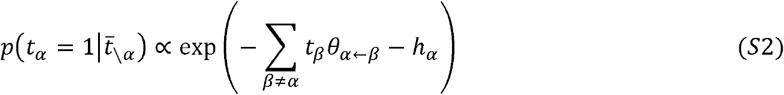

where *θ*_*α* ← *β*_ represents the effective coupling from *t*_*β*_ to *t*_*α*_ and *h*_*α*_ is the TF-specific bias term.

Importantly, we assume that TFs or TGs to regulate themselves, TGs are not permitted to regulate TFs, and we allow for asymmetric coupling (directed regulation) between TFs.

### 1.3 Model training

The model was trained using the scRNA-seq data described above, with parameters optimized to reproduce observed patterns of gene expression. Training employs gradient descent, with regularization applied to ensure biologically plausible network structures (no self-regulation, sparse interactions between TFs and TGs).

Let us denote by ***t***_*n*_ and ***g***_*n*_ the vectors representing binary activities of TFs and TGs in the n^th^ cell. We write

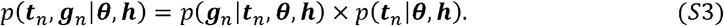

Let us define

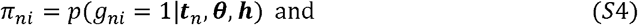

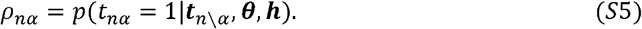

We have the total log-likelihood

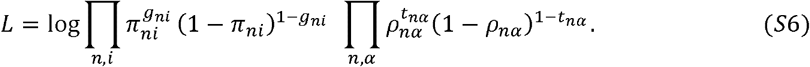

From this, we obtain the gradients

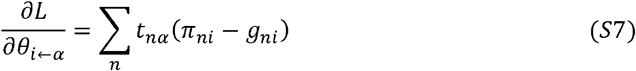

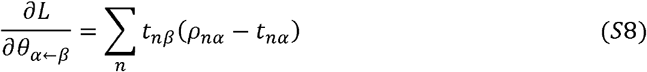

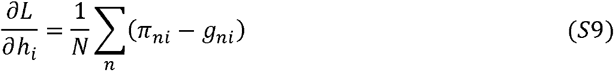

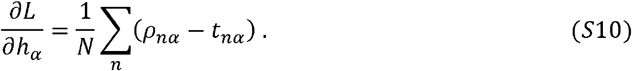

We regularize the inference using lasso regularization on *θ* s with a strength *λ* to make the network sparse. To pick the regularization strength, we employed a search over *λ* values, training our model on TFs and TGs and analyzing the accuracy of the statistics captured. To that end, we first set to zero interaction terms *θ*s whose magnitude was under 0.01 to zero. Using Markov chain Monte Carlo using Eq. S2 for TFs S1 for TGs, we generated samples to investigate predicted mean activities and correlations. Sparsity of individual models for all tissues are recorded in Table S1. For each model, we chose *λ* that led to the highest Pearson correlation coefficient for TF-TF correlations. Other statistics were analyzed to ensure reasonable fit over all (Fig S1-S10). The search over *λ* was performed over integer values. If no models had a good fit, we modified the search to a resolution of 0.1 near the best value. The values chosen are listed in Table S2.

We used the Adam Optimization Algorithm, an adaptive learning algorithm to speed up training while not missing areas of high likelihood. This tradeoff is regulated with what are called momentum and RMSprop. the standard parameters were used: *β*_1_ =0.9, *β*_2_ = 0.999, *α* =0.1, *ϵ* = 10^−8^. We ran the models for 10 iterations for each choice of *λ* and chose the one with the highest likelihood for further analysis. Finally, we note that for each tissue we learnt a separate model for TFs and TGs for the ages of 3m and 24m.

## Section 2. Information theoretic quantification

### 2.1 Computing mutual information

Mutual information between all TFs and a given TG is expressed as follows:

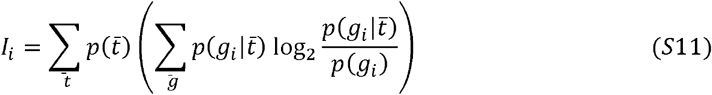

Where 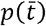 is the stationary distribution of TFs, generated using Markov Chain Monte Carlo using Eq. S2. Specifically, first, a set of TF activities are randomly sampled from the data. In each iteration, a TF is randomly chosen and is activated with a probability

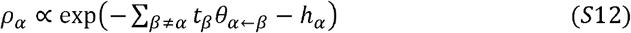

and deactivated with probability 1 − *ρ*_*α*_ The Markov chain is run for a burn-in period of 200000 iterations. Then it is sampled every 1000 iterations and saved until a total of 60000 samples are obtained across multiple chains. We then use these samples to calculate gene activation probabilities 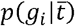 using equation S1 and to find the marginal probabilities 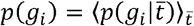.

We complete this whole process 10 times and averaged the MI for each gene over the 10 calculations. This calculation was performed separately for models trained on data from young mice (3m) and old mice (24m) to analyze the difference in the MI values for each age.

### 2.2 Estimating the effecting of rejuvenation

To estimate the *in silico* effect of rejuvenation, we performed the heterochronic MI calculation by substituting the stationary distribution of TFs in old mice with the one from young mice and then again by replacing the gene regulatory network in the old mice with that of the young mice to identify the causes of the decrease in MI with age. (Figs S11-S20).

## Section 3. Structural analysis of regulatory networks

### 3.1 Quantifying structural changes in the TF-TF network

To look at the structure of the TF-TF network, we first analyzed the out-degree distribution. Out degree of a TF node is the total number of other TFs that a particular TF controls. Prior to structural analysis, we recognized that the optimal density of interactions was different between ages. To remove confounding factors related to edge density, in any comparison, we rarified the denser network to the less dense network by randomly removing edges. This calculation was performed 10 times. We also include a null model, where a network of the same size as the others is randomly populated with connections until it has an equal number of connections. This calculation is also repeated 10 times. After these controls, we found that the distribution of out degrees for 24m networks almost always has a longer tail than the out-degree distribution for the 3m network. Notably, both networks have wider degree distribution spread compared to the null model (see figures S21-S30). This suggests that the networks in 24m had more hubs (fewer TFs regulating more genes) compared to the 3m networks.

We next evaluated the number of adapting loops in the network. We consider two types of loops that are known to perform adaptation: incoherent feedforward (IFFL) and incoherent feedback (IFL). The primary example of IFFL is one in which a regulator activates both a target and its inhibitor. The IFL is a regulator activating a target and the target activates its own inhibitor. There are other realizations of these loops, pictured in the diagrams in figures S31 and S32. We wanted to quantify whether cells from older organisms lose the ability to adapt to random perturbations. Notably, the total count for each loop type (IFL and IFFL) was significantly lower in 24m (Table S4, Figure S33-S43) across all tissues.

### 3.2 Stability to perturbations

After we found that cells from older mice have a disproportionately lower number of adapting loops in the TF-TF network, we decided to test the networks’ stability to random perturbations. To accomplish this, we randomly chose 10% of TFs in a tissue. If the average activity of the TF 50%, we performed a knock in (*p*(*t*_*α*_ = 1) = 1). In contrast, if the average activity was < 50%, we performed a knock-out (*p*(t_*α*_ = 1) = 0). With these constraints on some of the TFs, we sampled the rest of the TF network and predicted activity change of the rest of the TFs. We found that the activity expression in the perturbed state was much closer to the original state in young mice than in old mice in all tissues except the thymus, where the change was slightly higher in 3m. This was quantified with the L1 norm of TF activity. For each age, we quantified 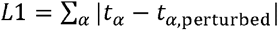, normalized by the total average TF activity for that age (see Figure S44).

### 3.3 Computational knock-ins

To identify master regulator TFs, we performed in-silico knock-in (KI) experiments. To focus our attention, we studied cells from limb muscles. Our task was to identify how knock-in of a subset of TFs changes information flow in the GRN. This is how the calculation was performed for each TF. First, we used MCMC on models learnt on cells from old mice to generate TF samples. In the MCMC, we knocked in the designated TF to be always active, thereby biasing the entire distribution. We computed the MI between genes and TFs in this perturbed sample. We performed this calculation for each TF and computed the L1 norm of MI between young cells and the perturbed old cells. To identify the combined effect of simultaneous activation of multiple TFs, we repeated this calculation by simultaneously turning on top 5 TFs.

