## Supplementary Tables and Figures for "Decay in transcriptional information flow is a hallmark of cellular aging"

### 1. Tables

**Table S1. Optimal sparsity of inferred gene regulatory networks**

| | TF | | TG | |
|---|---|---|---|---|
| **Tissue** | 3m | 24m | 3m | 24m |
| GAT | 0.371 | 0.157 | 0.361 | 0.151 |
| Limb Muscle | 0.180 | 0.056 | 0.175 | 0.053 |
| Lung | 0.207 | 0.064 | 0.205 | 0.067 |
| Trachea | 0.239 | 0.123 | 0.234 | 0.123 |
| SCAT | 0.354 | 0.176 | 0.337 | 0.172 |
| Pancreas | 0.237 | 0.162 | 0.231 | 0.158 |
| Thymus | 0.128 | 0.093 | 0.135 | 0.097 |
| Tongue | 0.211 | 0.170 | 0.205 | 0.167 |
| Large Intestine | 0.163 | 0.139 | 0.156 | 0.134 |
| Skin | 0.253 | 0.160 | 0.246 | 0.153 |

**Table S2. Optimal regularization parameter $\lambda$**

| Tissue | 3m | 24m |
|---|---|---|
| GAT | 0.98 | 2 |
| Limb Muscle | 2 | 3 |
| Lung | 2 | 5 |
| Trachea | 2 | 2 |
| SCAT | 1 | 1.3 |
| Pancreas | 3 | 2 |
| Thymus | 2 | 3 |
| Tongue | 3 | 3 |
| Large Intestine | 5 | 3 |
| Skin | 3 | 2 |

**Table S3. Mean MI (bits)**

| | | | Rejuvenated | |
|---|---|---|---|---|
| Tissue | 3m | 24m | Distribution | Network |
| GAT | 0.234 | 0.054 | 0.171 | 0.0925 |
| Limb Muscle | 0.099 | 0.020 | 0.062 | 0.0364 |
| Lung | 0.065 | 0.026 | 0.032 | 0.0534 |
| Trachea | 0.080 | 0.061 | 0.079 | 0.0587 |
| SCAT | 0.256 | 0.095 | 0.229 | 0.0986 |
| Pancreas | 0.132 | 0.076 | 0.149 | 0.0574 |
| Thymus | 0.066 | 0.051 | 0.049 | 0.0644 |
| Tongue | 0.100 | 0.088 | 0.101 | 0.0868 |
| Large Intestine | 0.100 | 0.091 | 0.118 | 0.0737 |
| Skin | 0.083 | 0.089 | 0.125 | 0.0573 |

**Table S4. Loop Counts**

| | IFL Count $\times 10^{-4}$ | | | IFFL Count $\times 10^{-4}$ | | |
|---|---|---|---|---|---|---|
| Tissue | 3m | 24m | $\Delta Count$ | 3m | 24m | $\Delta Count$ |
| GAT | 118.35 | 101.37 | 16.98 | 74.23 | 6.07 | 68.16 |
| Limb Muscle | 17.47 | 15.79 | 1.68 | 8.15 | 0.37 | 7.78 |
| Lung | 22.33 | 19.93 | 2.41 | 11.67 | 0.45 | 11.22 |
| Trachea | 100.96 | 80.99 | 19.98 | 39.10 | 4.73 | 34.37 |
| SCAT | 162.65 | 138.24 | 24.41 | 93.33 | 12.58 | 80.75 |
| Pancreas | 131.34 | 104.89 | 26.45 | 28.25 | 6.28 | 21.97 |
| Thymus | 70.74 | 50.23 | 20.51 | 13.63 | 1.71 | 11.92 |
| Tongue | 125.94 | 94.13 | 31.82 | 16.34 | 3.22 | 13.12 |
| Large Intestine | 131.69 | 112.14 | 19.55 | 10.37 | 4.26 | 6.12 |
| Skin | 126.22 | 90.38 | 35.84 | 33.56 | 4.49 | 29.07 |

#### 2. Figures

##### A. Model Statistics of Tissues.

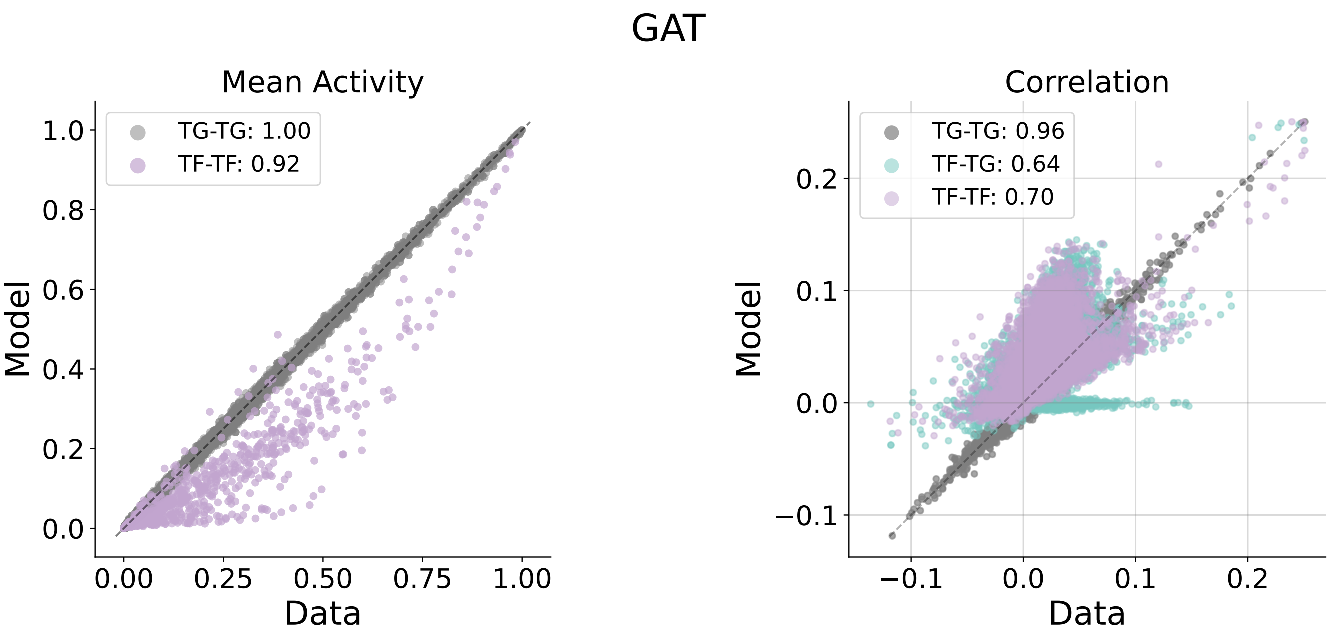


**3m**

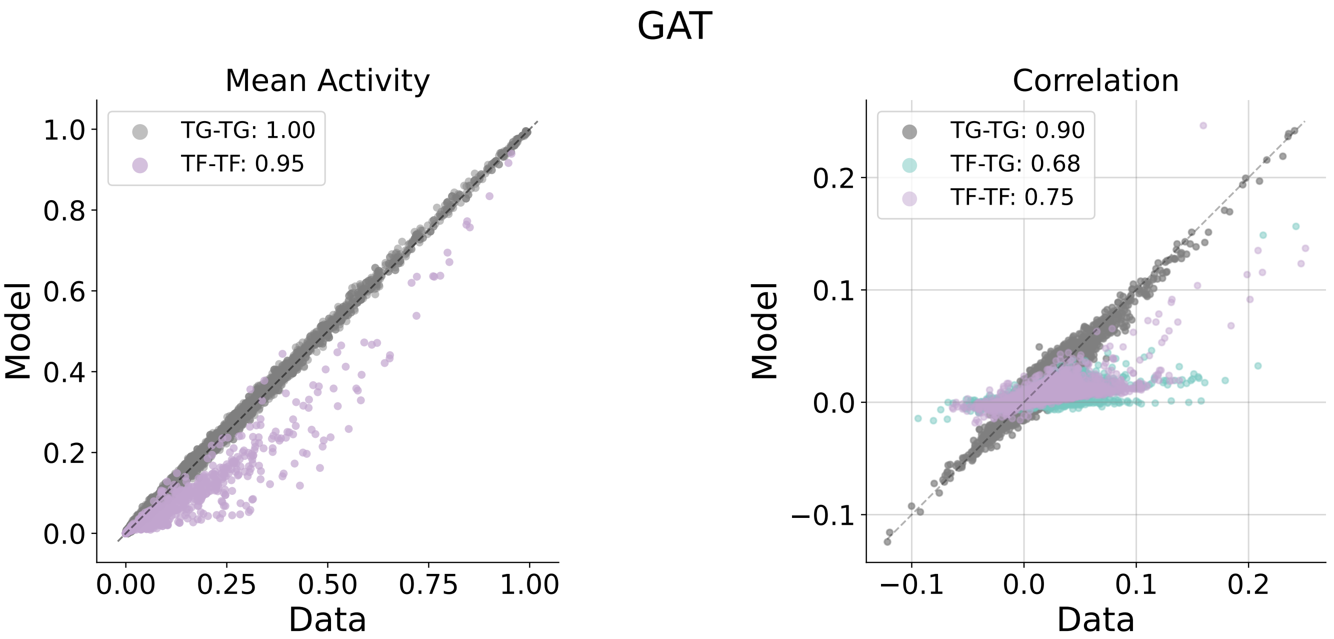


**24m**

Fig. S1. **GAT statistics across 3m and 24m.**

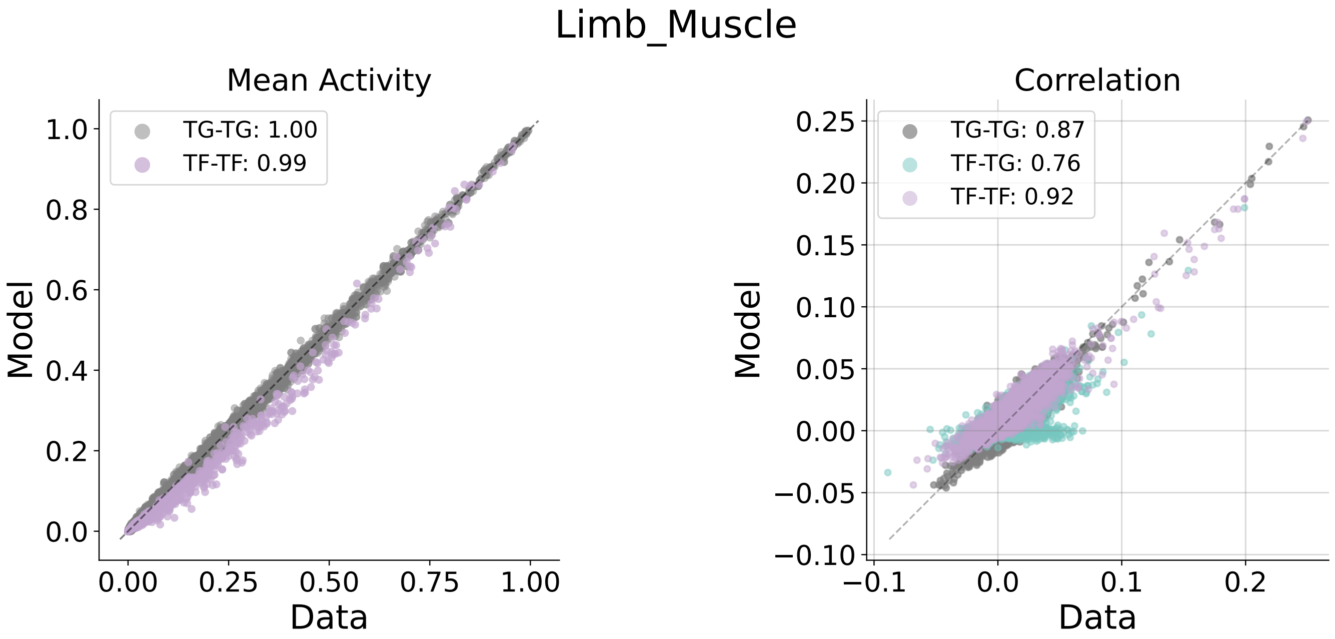


**3m**

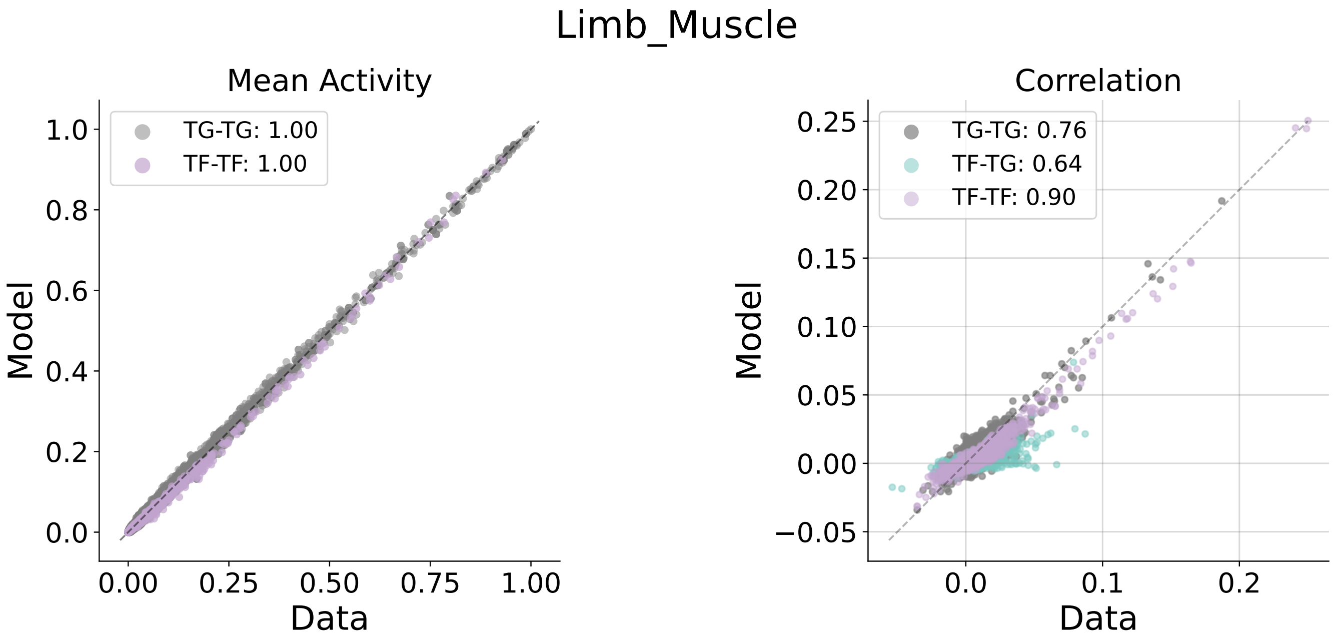


**24m**

Fig. S2. **Limb Muscle statistics across 3m and 24m.**

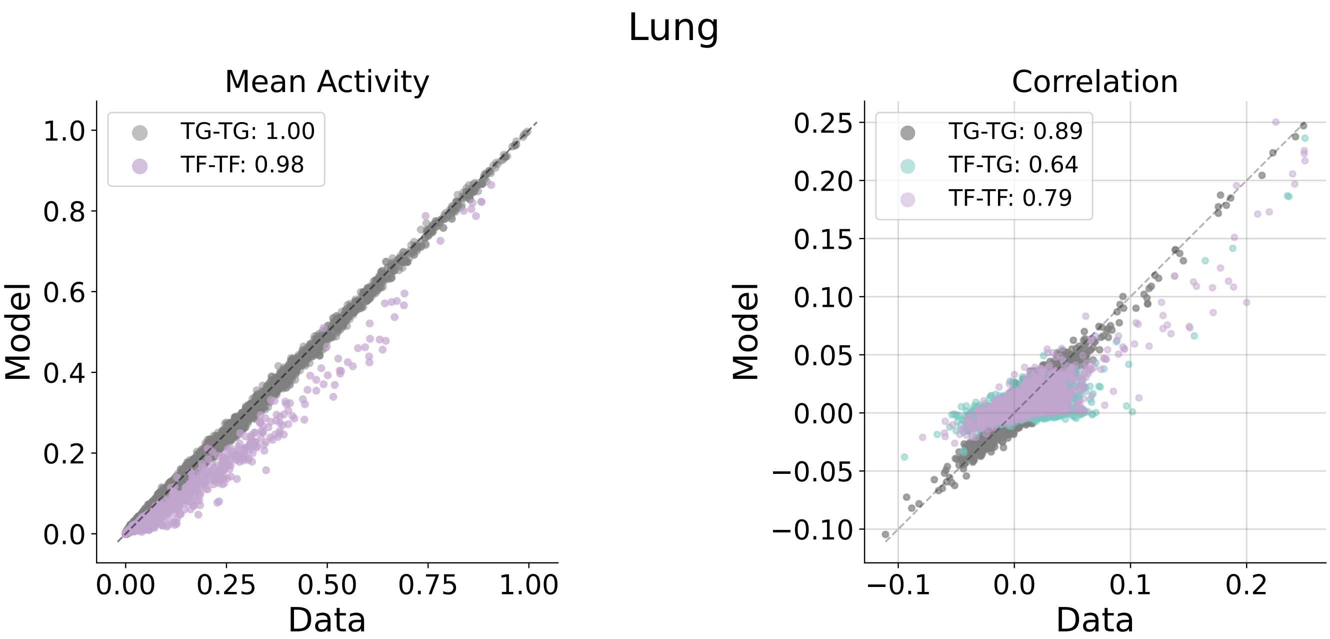


**3m**

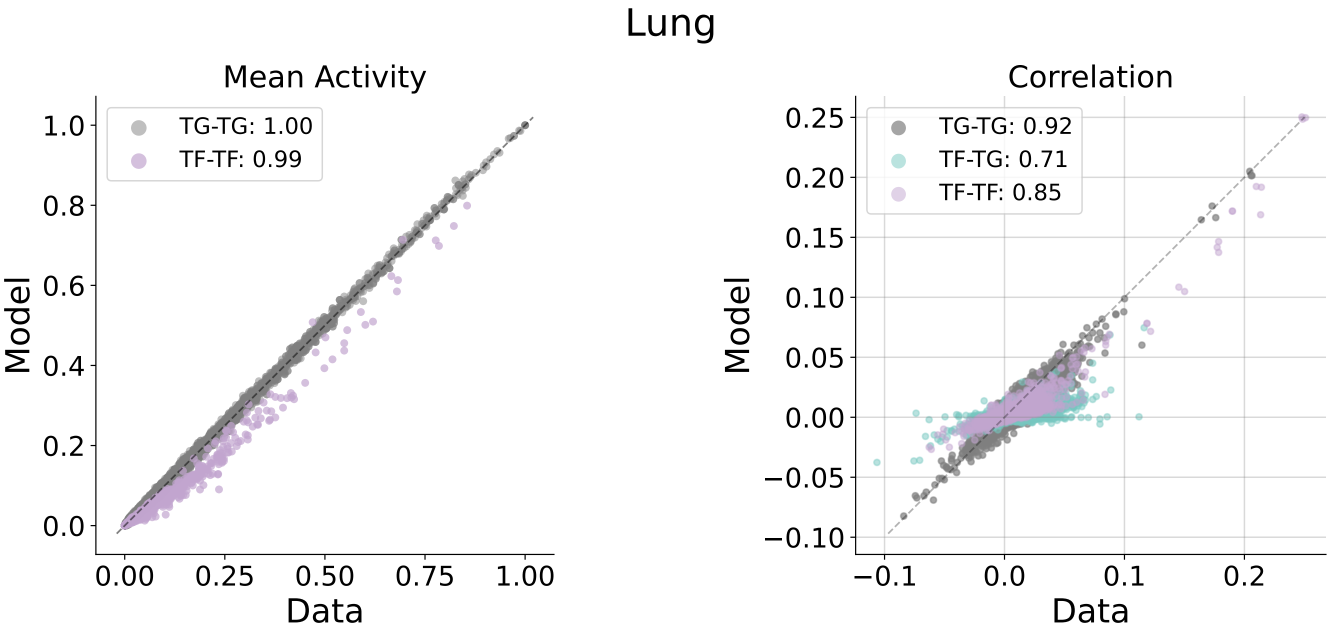


**24m**

Fig. S3. **Lung statistics across 3m and 24m.**

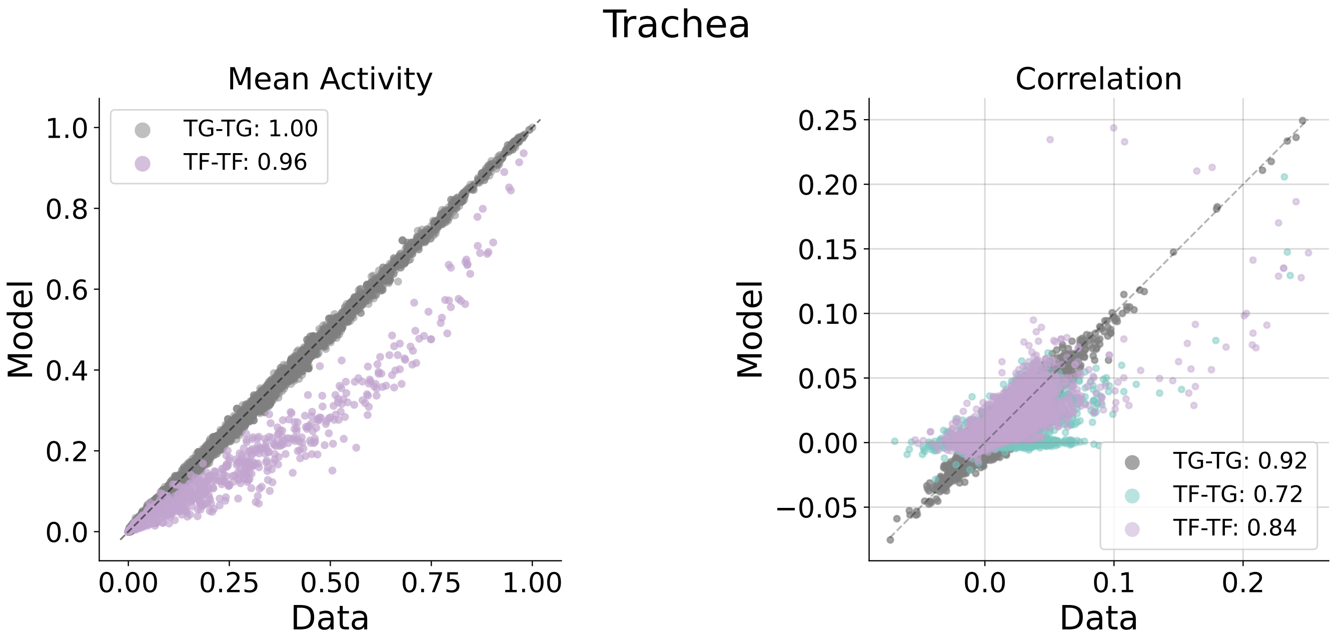


**3m**

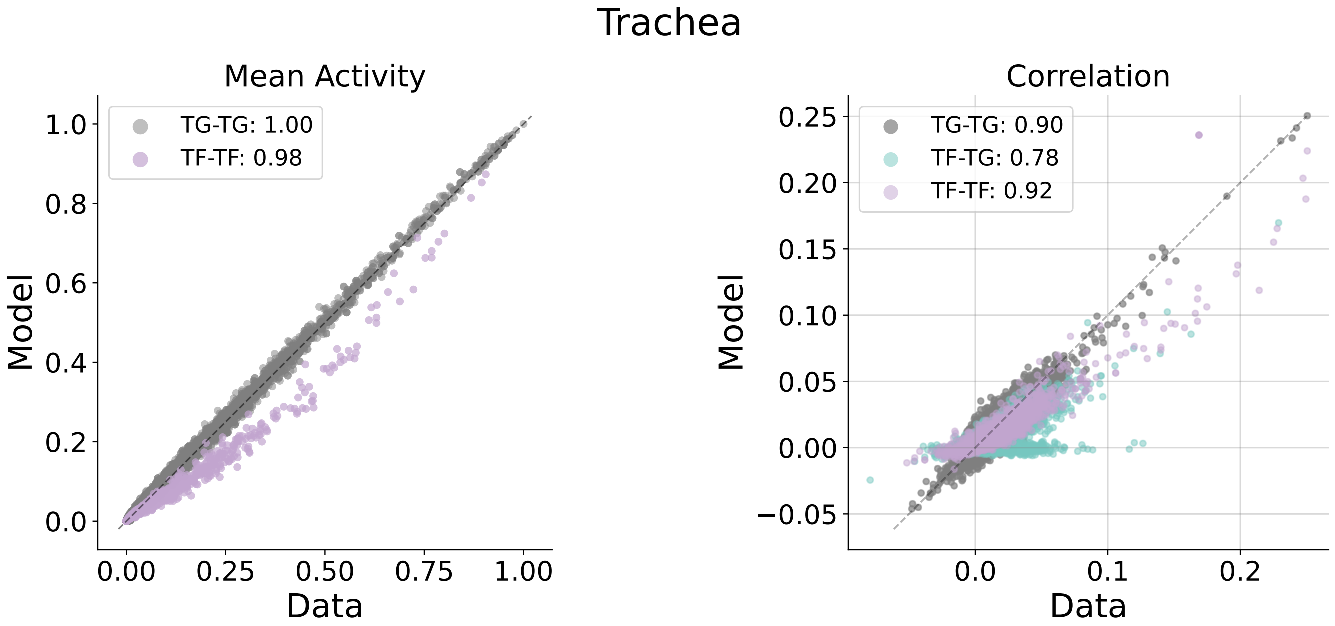


**24m**

**Fig. S4. Trachea statistics across 3m and 24m.**

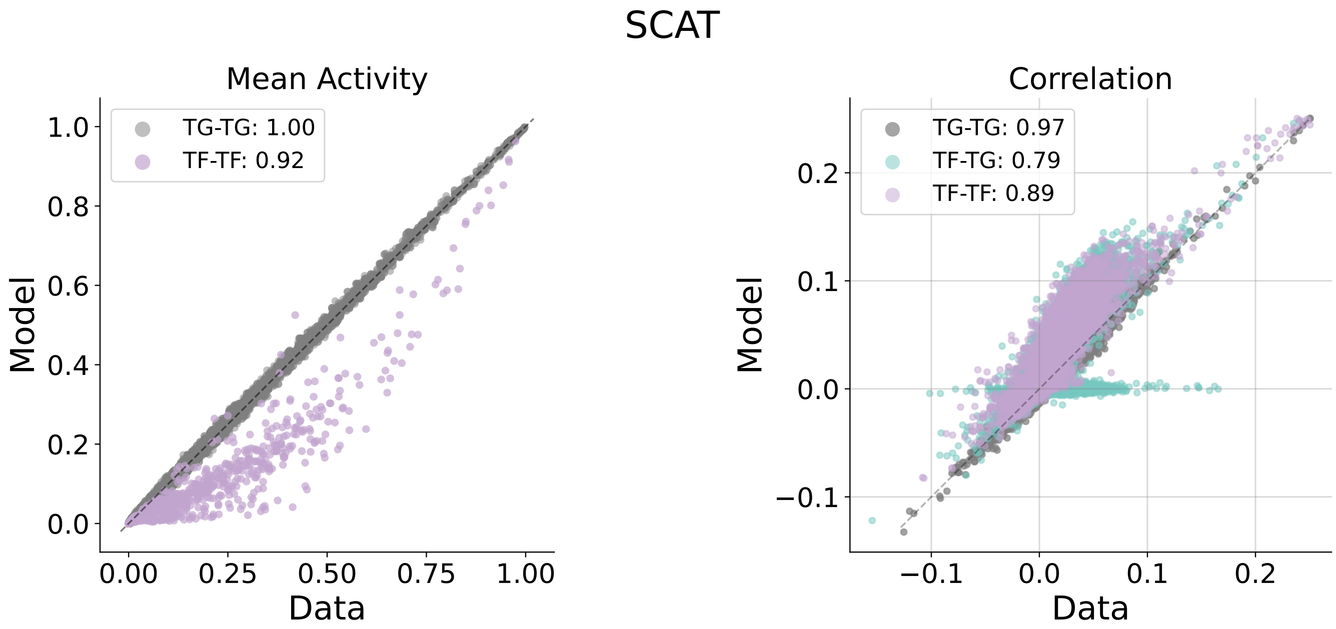


**3m**

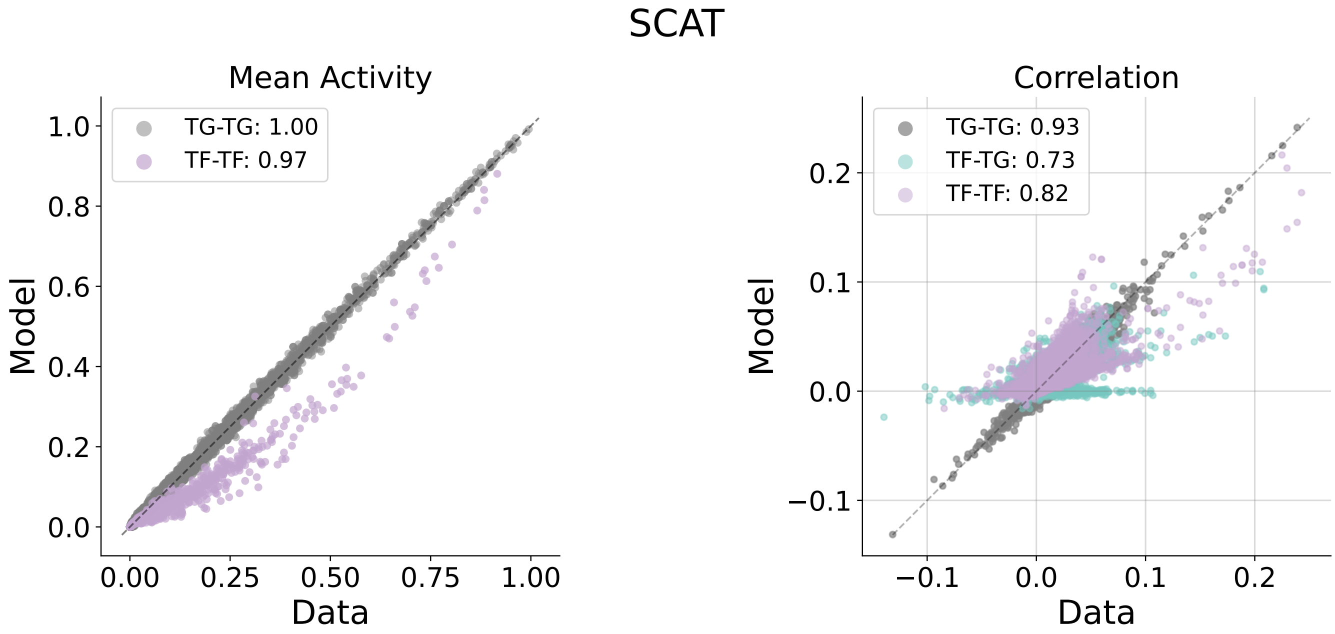


**24m**

**Fig. S5. SCAT statistics across 3m and 24m.**

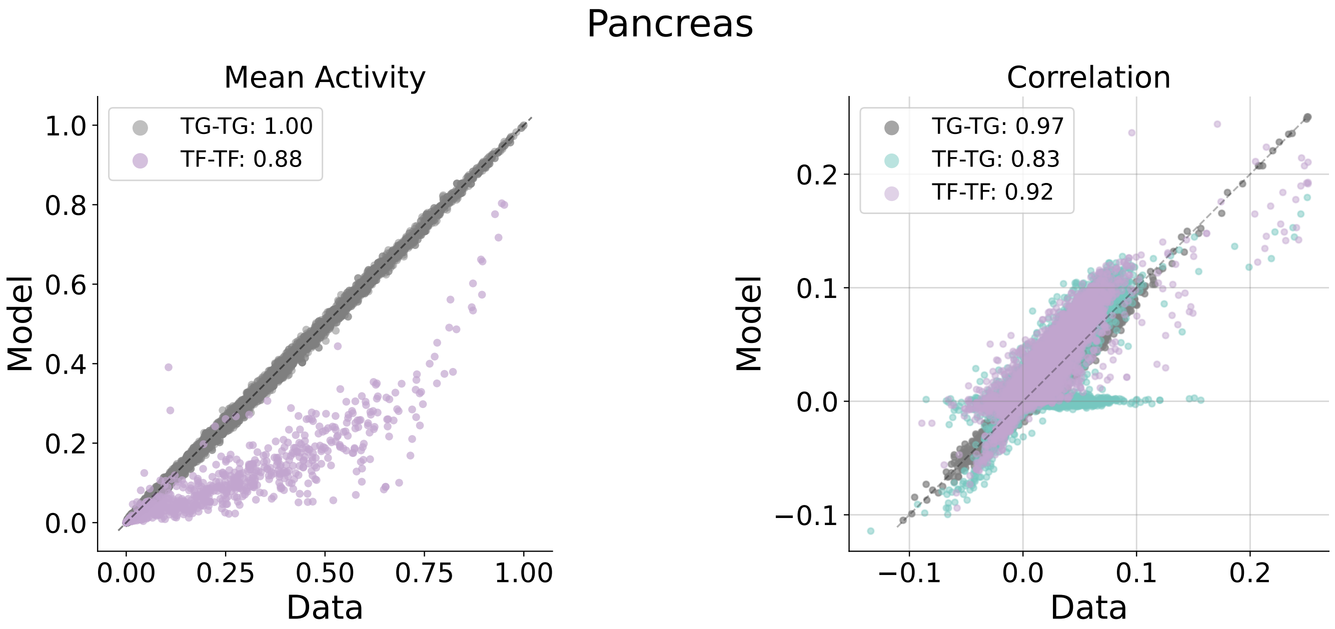


**3m**

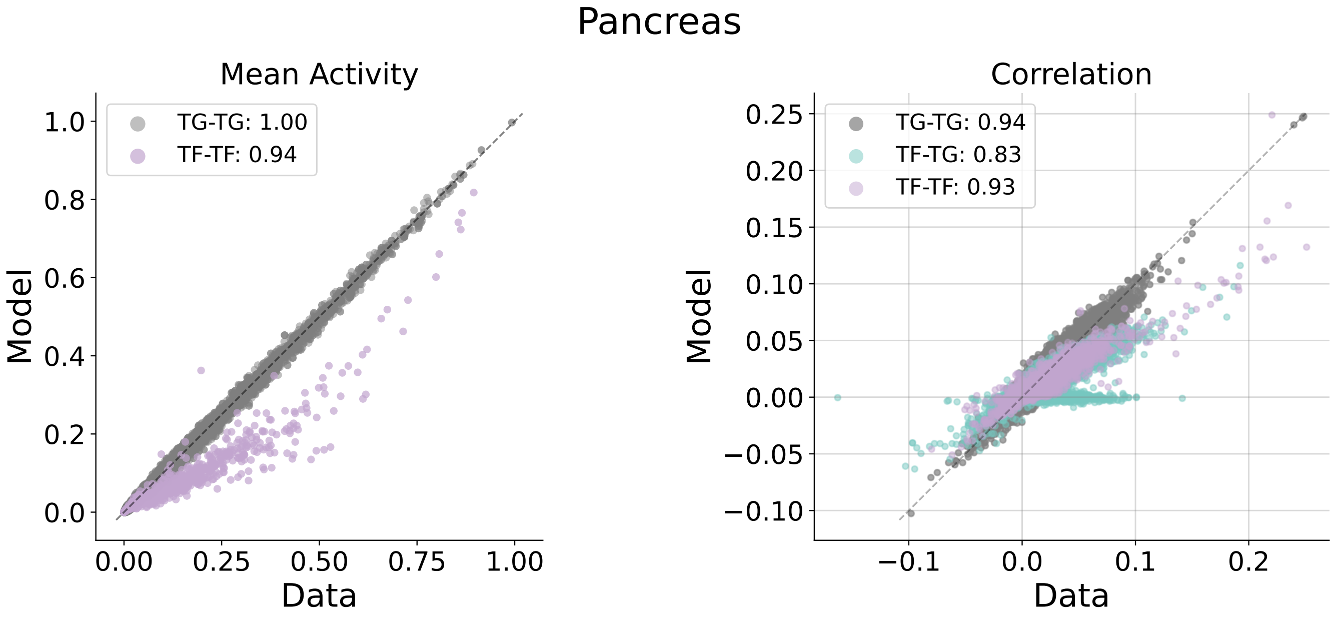


**24m**

Fig. S6. **Pancreas statistics across 3m and 24m.**

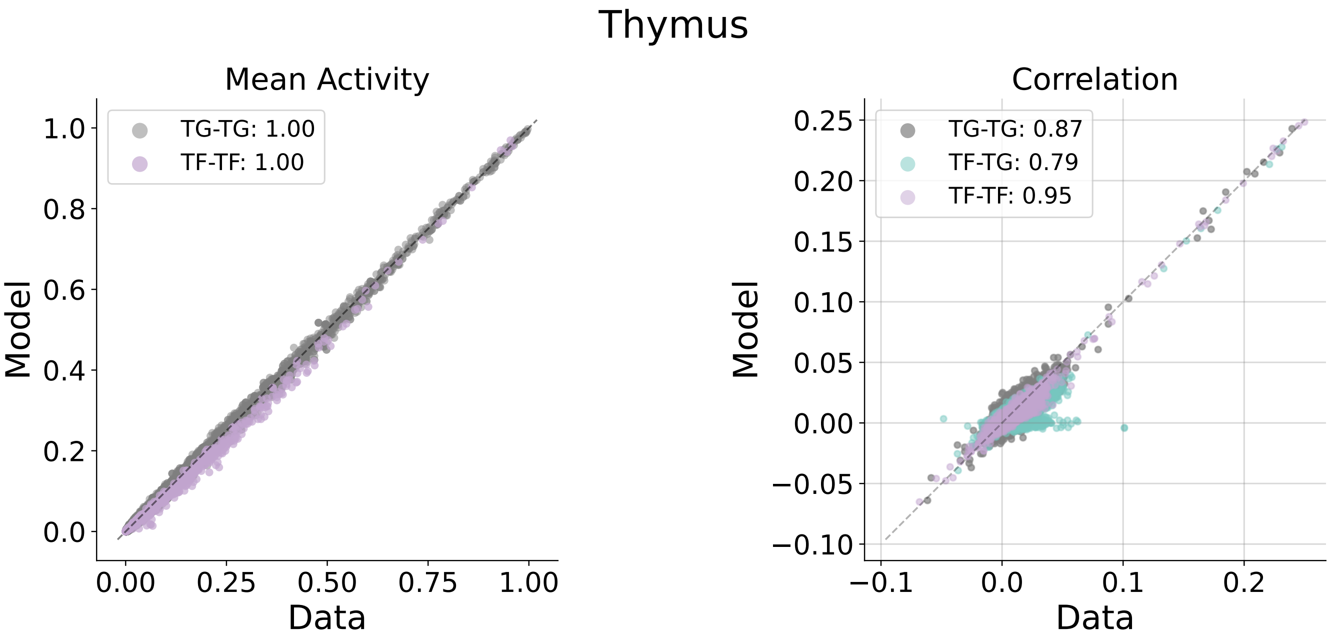


**3m**

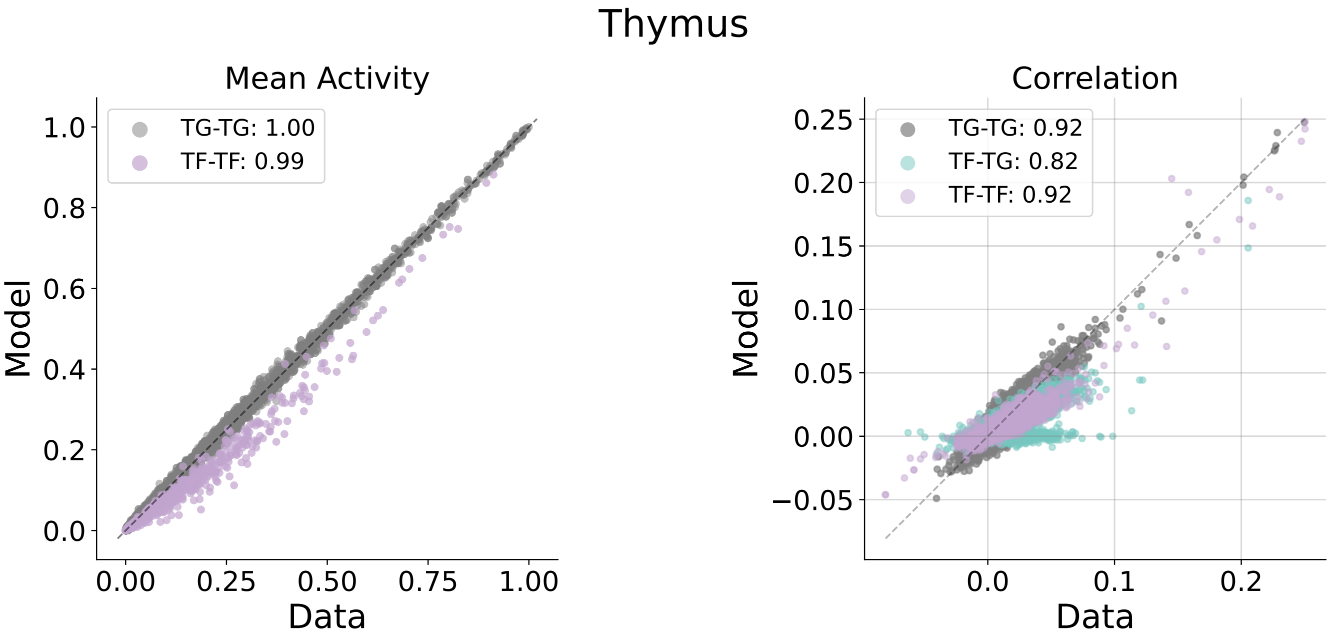


**24m**

Fig. S7. **Thymus statistics across 3m and 24m.**

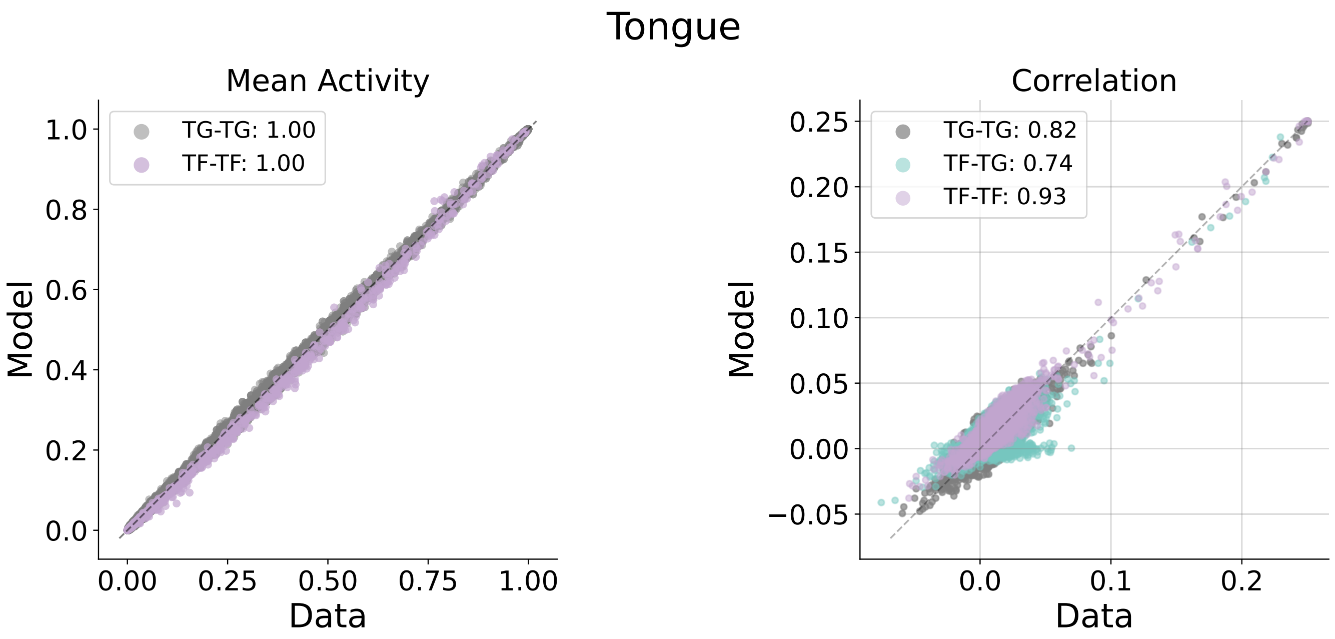


**3m**

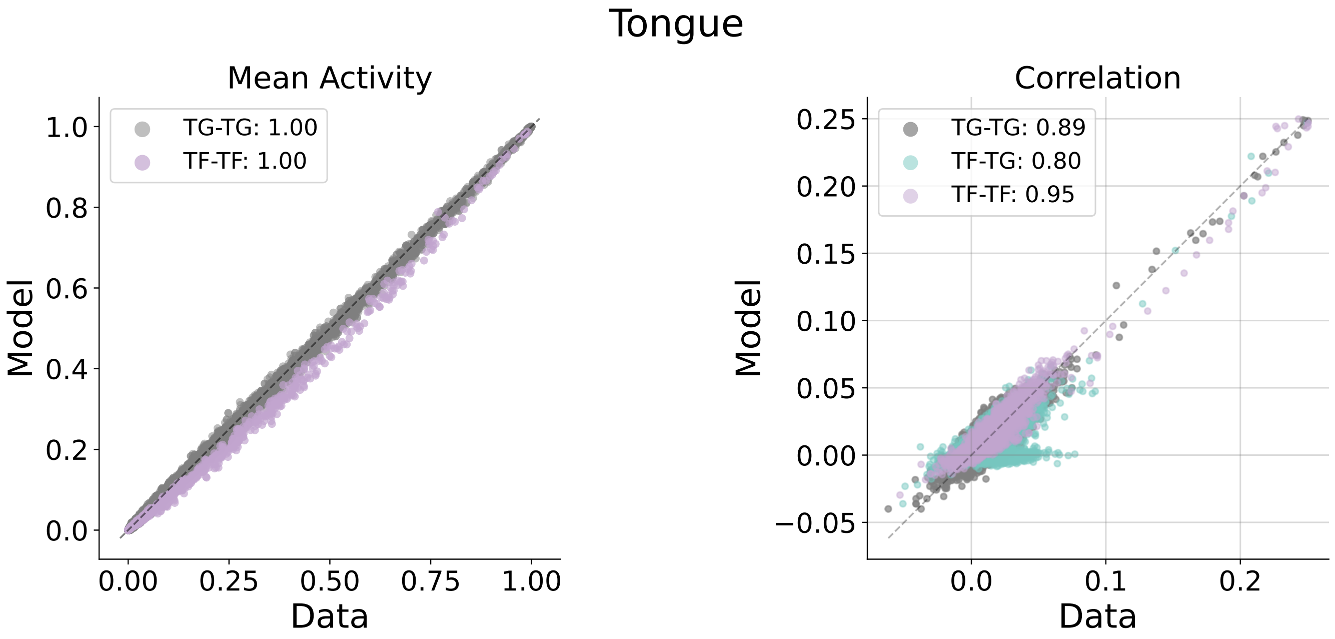


**24m**

Fig. S8. **Tongue statistics across 3m and 24m.**

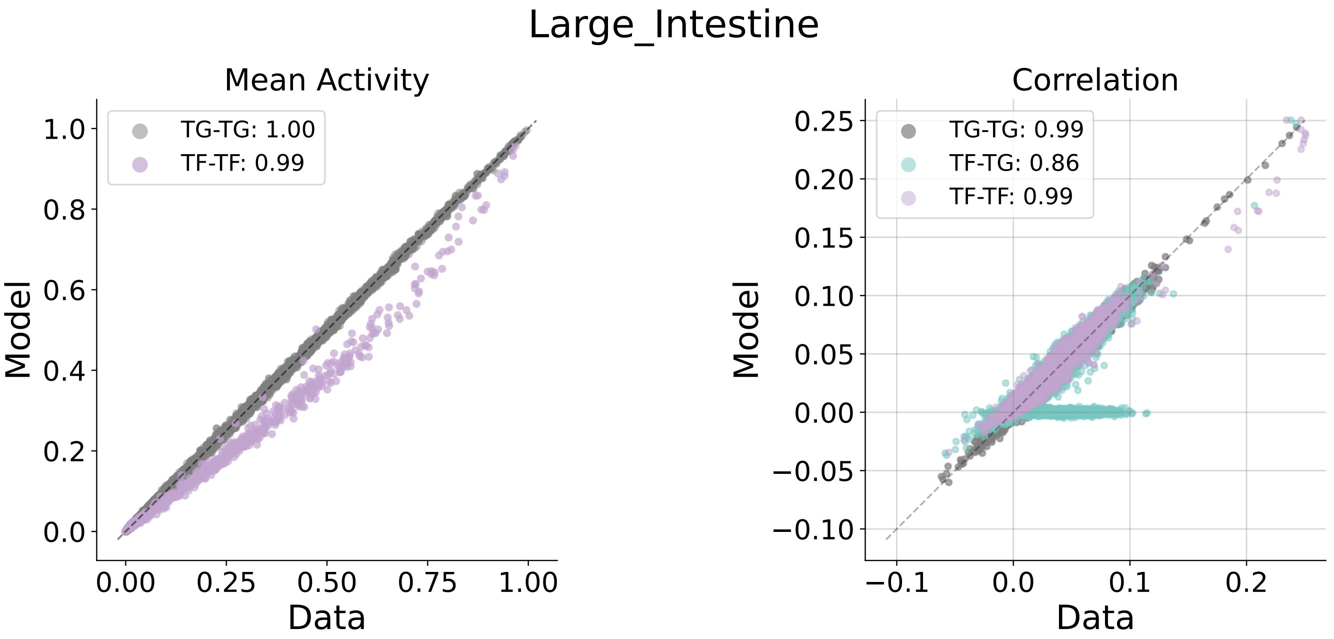


**3m**

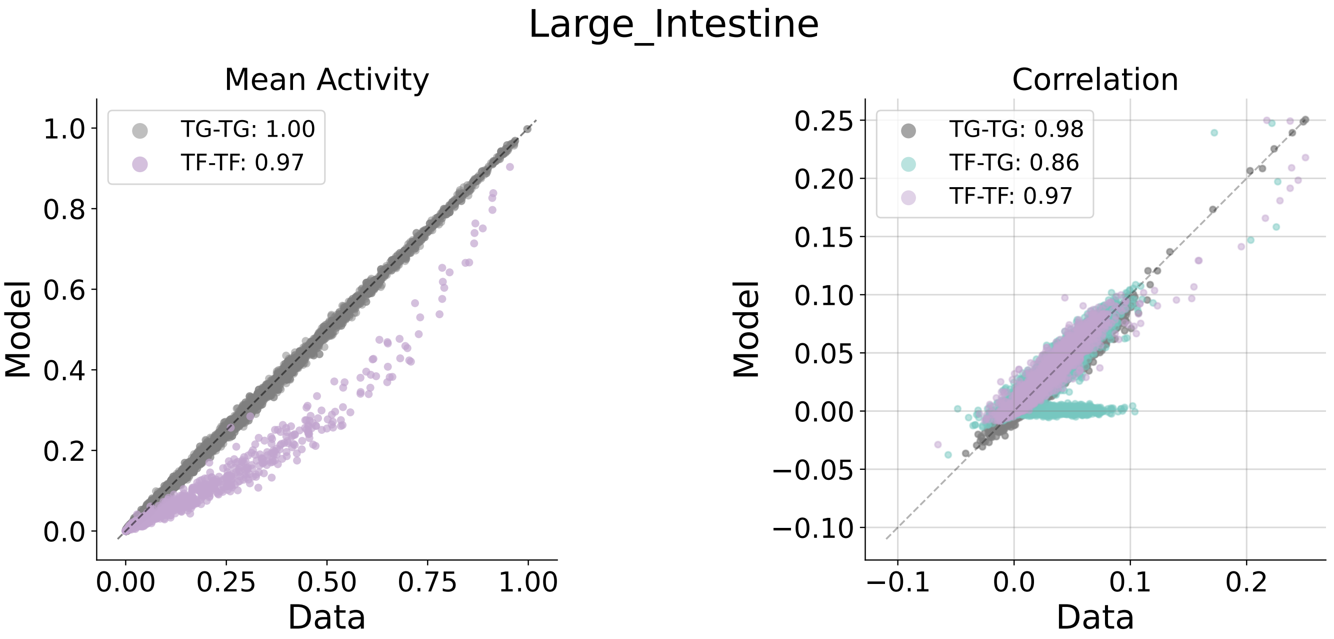


**24m**

Fig. S9. **Large Intestine statistics across 3m and 24m.**

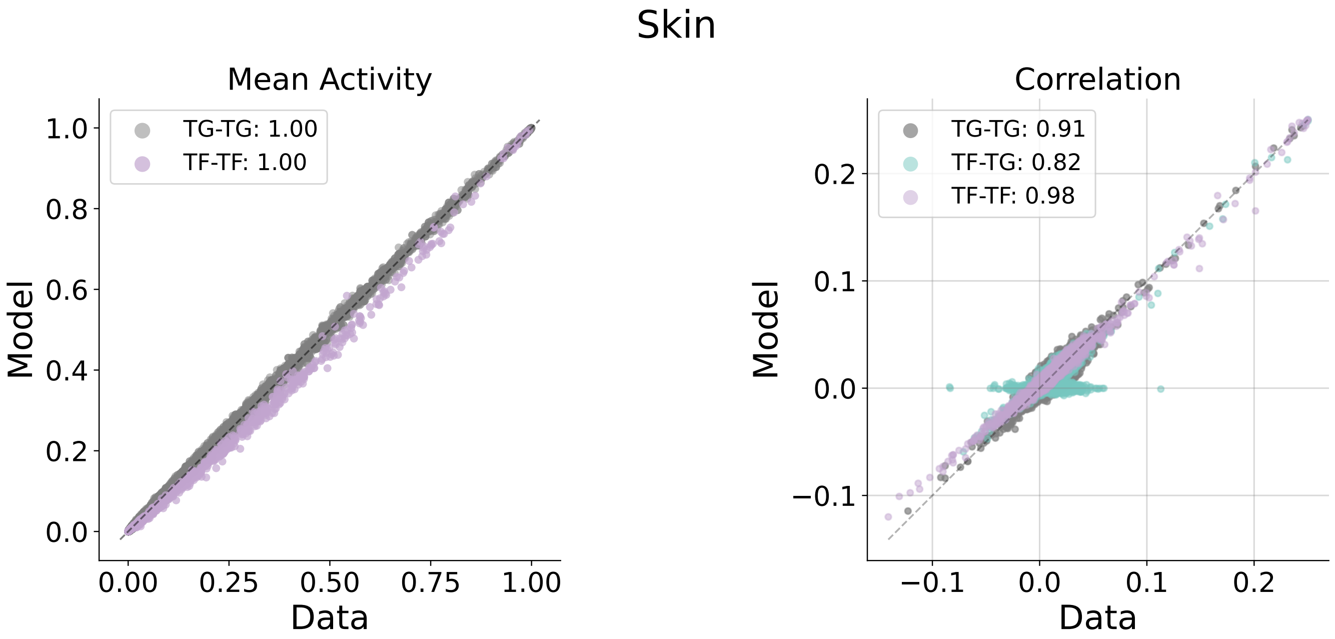


**3m**

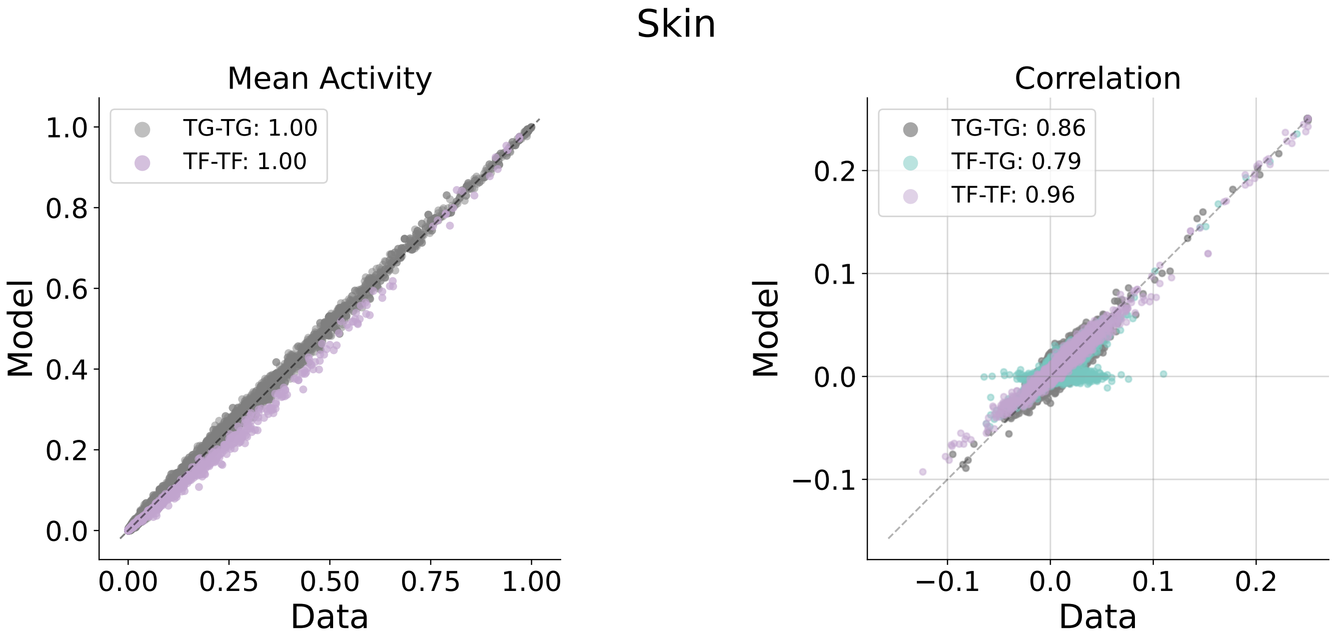


**24m**

**Fig. S10. Skin statistics across 3m and 24m.**

#### B. MI Swap Results.

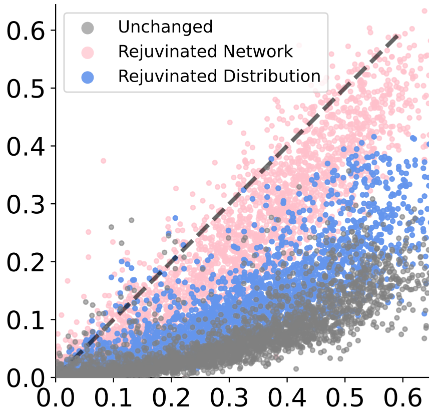


**Fig. S11. GAT.**

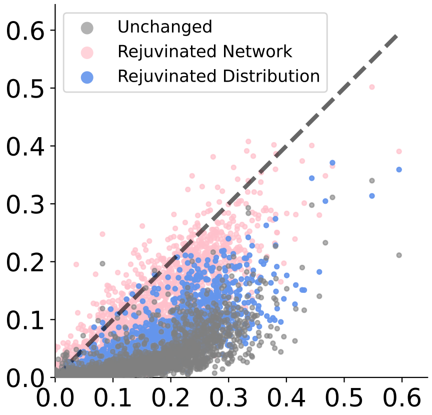


**Fig. S12. Limb Muscle.**

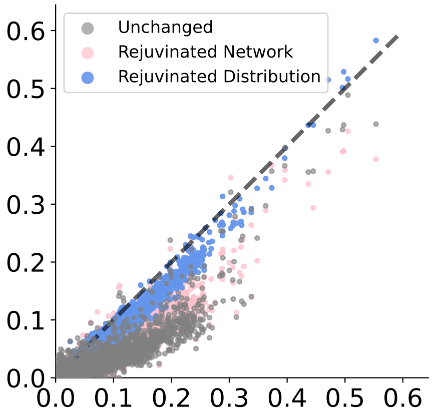


**Fig. S13. Lung.**

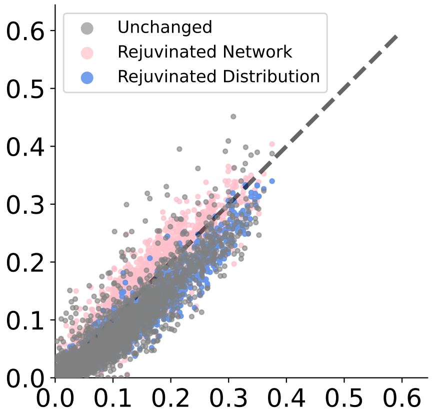


**Fig. S14. Trachea.**

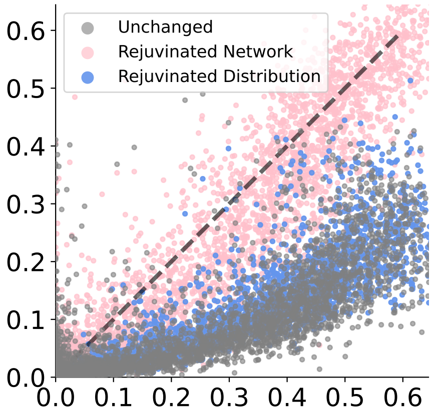


**Fig. S15. SCAT.**

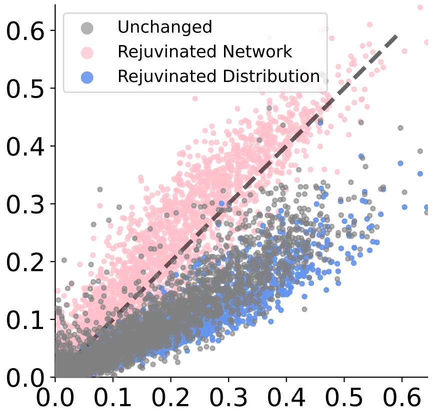


**Fig. S16. Pancreas.**

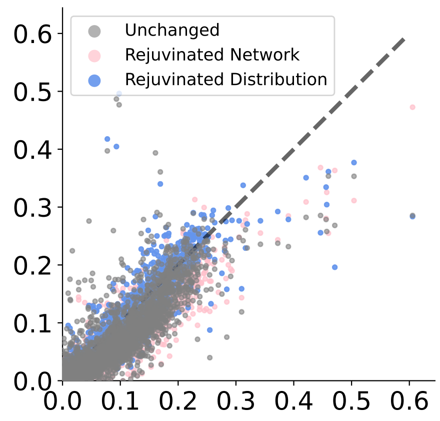


**Fig. S17. Thymus.**

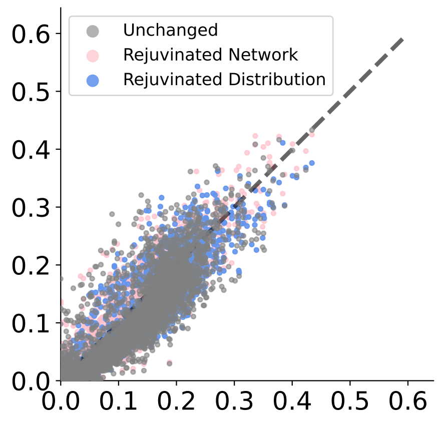


**Fig. S18. Tongue.**

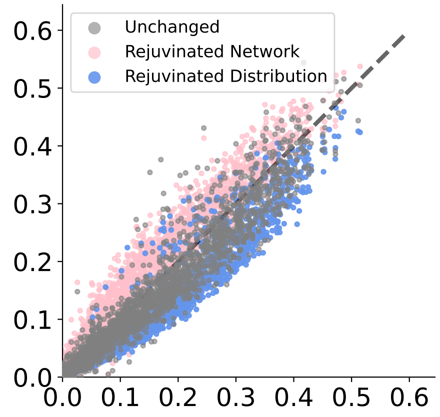


**Fig. S19. Large Intestine.**

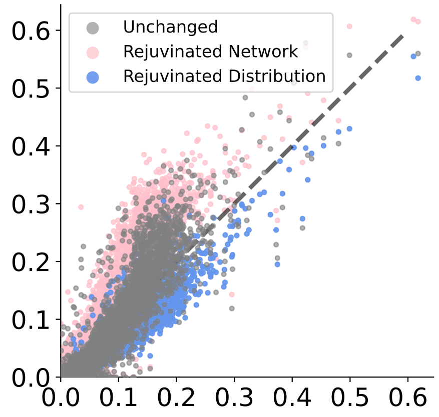


**Fig. S20. Skin 24m.**

#### C. Out Degrees TF Corrected.

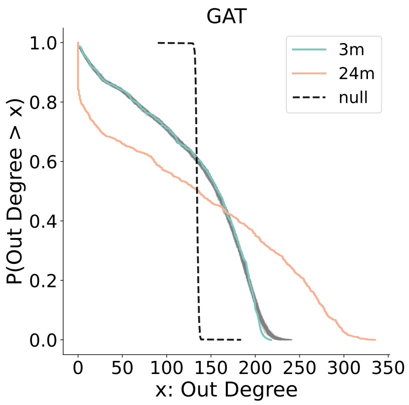


**Fig. S21. GAT.**

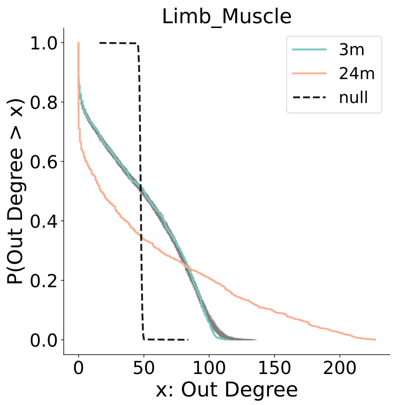


**Fig. S22. Limb Muscle.**

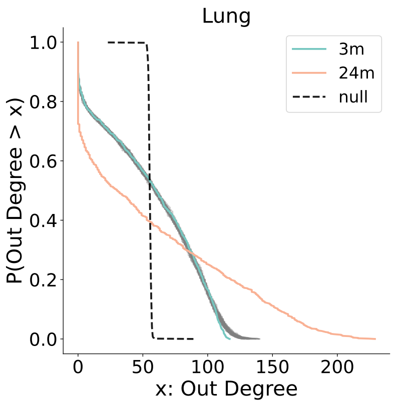


**Fig. S23. Lung.**

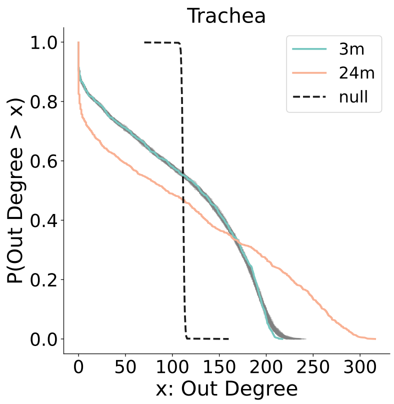


**Fig. S24. Trachea.**

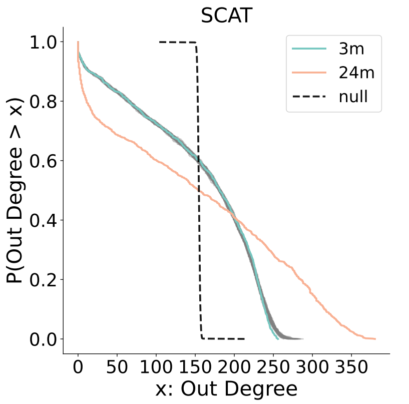


**Fig. S25. SCAT.**

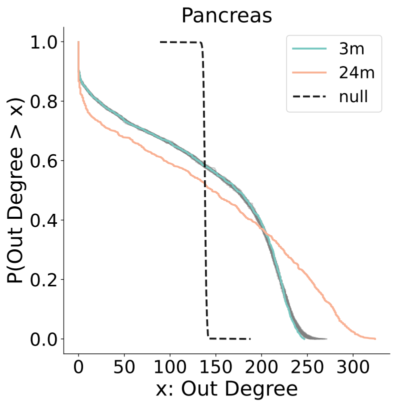


**Fig. S26. Pancreas.**

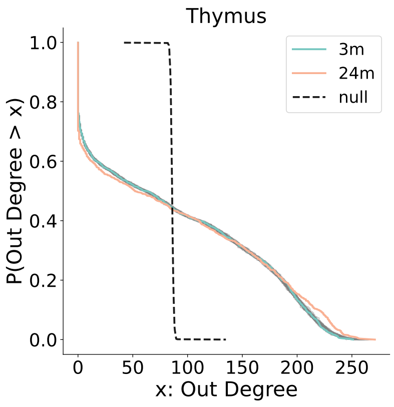


**Fig. S27. Thymus.**

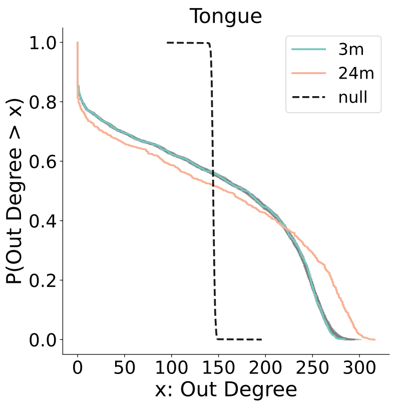


**Fig. S28. Tongue.**

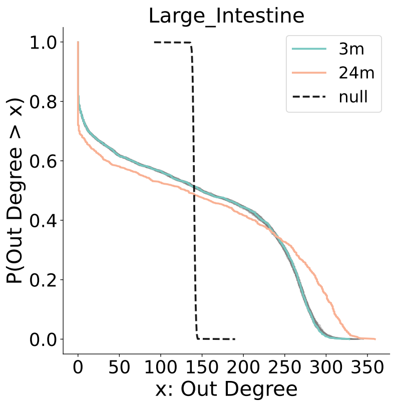


**Fig. S29. Large Intestine.**

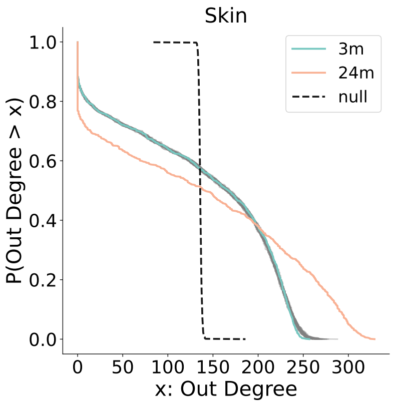


**Fig. S30. Skin 24m.**

#### D. Network Structure.

##### *D.1. Loops.*

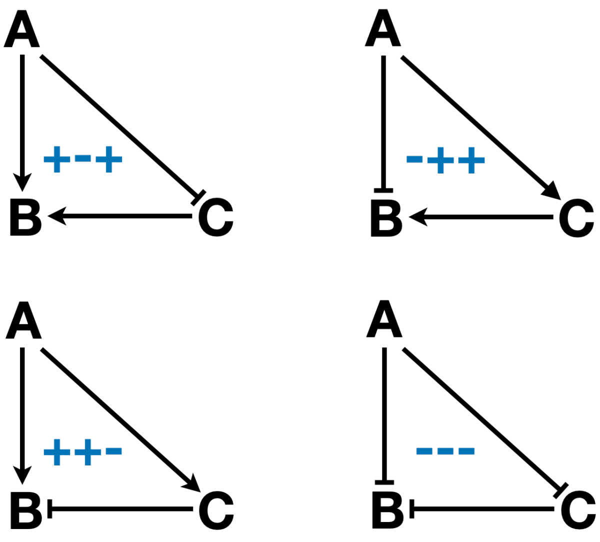


**Fig. S31.** Diagram of IFFL loop types.
"+": promotes, "-": inhibits

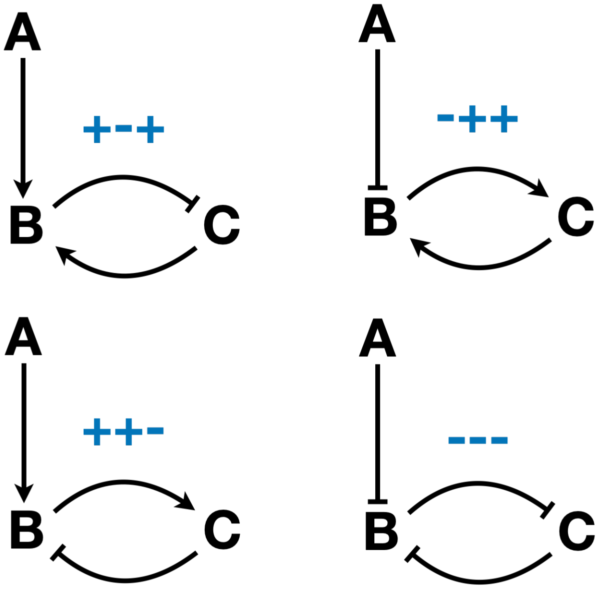


**Fig. S32.** Diagram of IFL loop types.
"+" : promotes, "-": inhibits

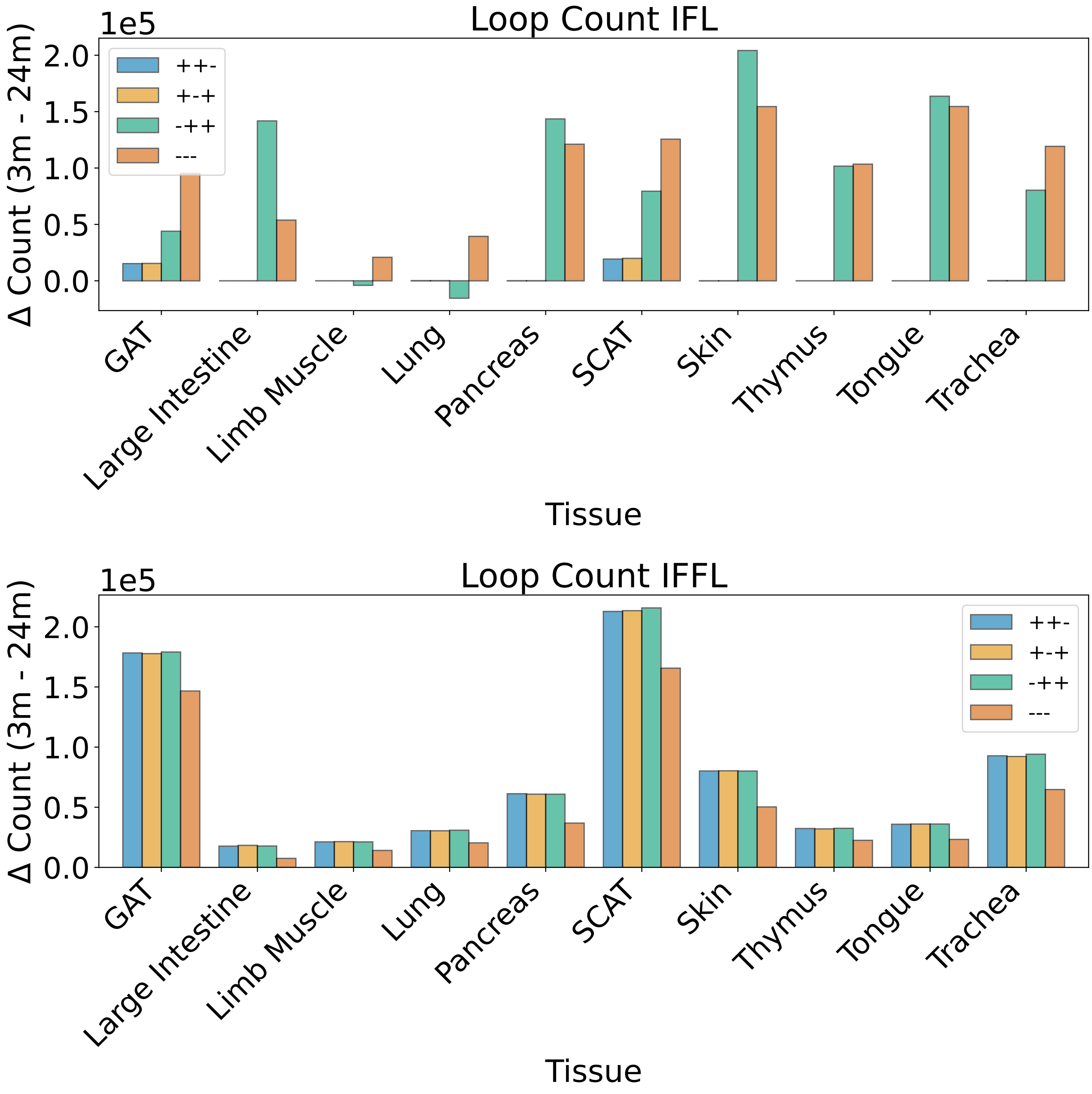


**Fig. S33.** Loop Counts Summary IFFL and IFL.

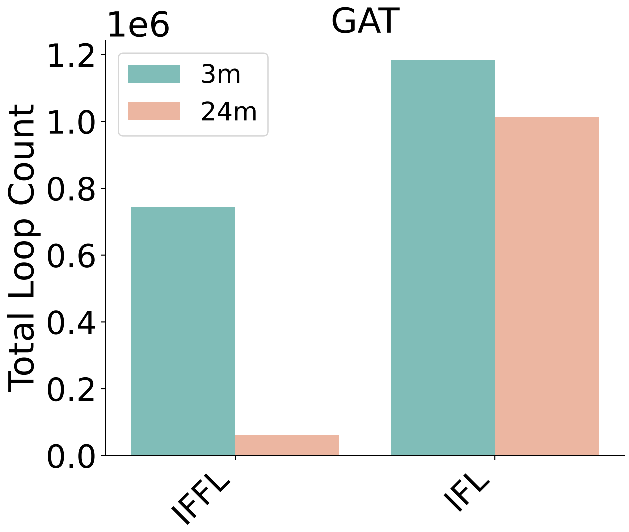


**Fig. S34. GAT.**

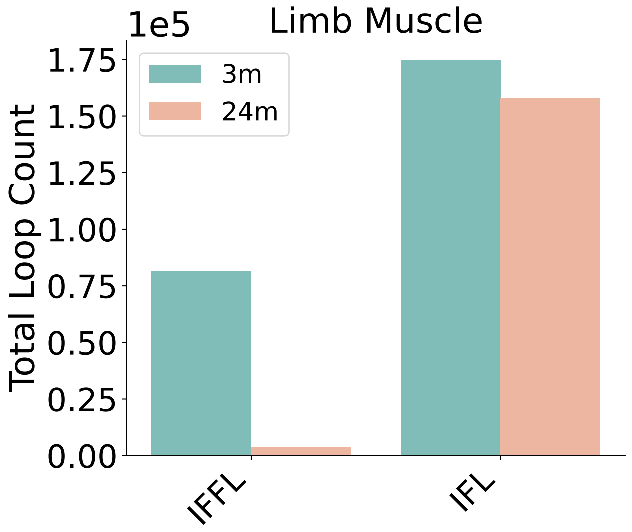


**Fig. S35. Limb Muscle.**

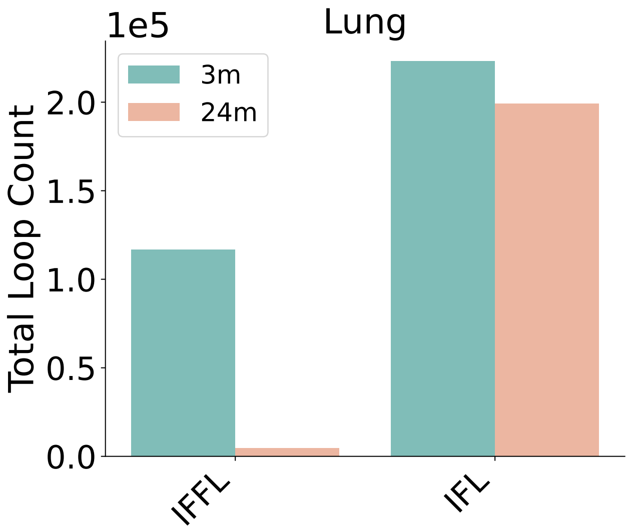


**Fig. S36. Lung.**

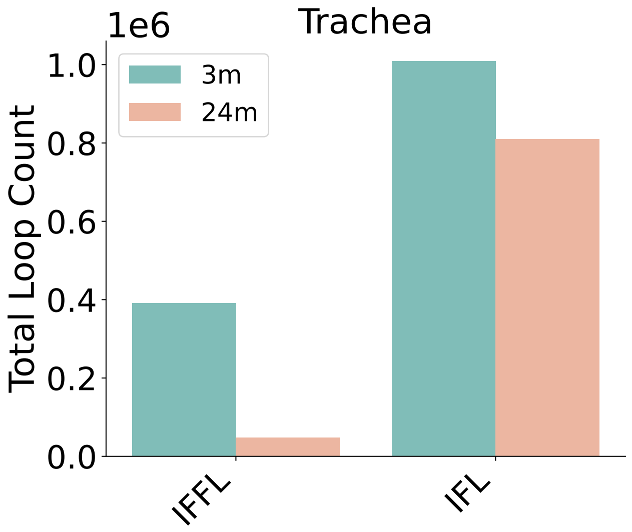


**Fig. S37. Trachea.**

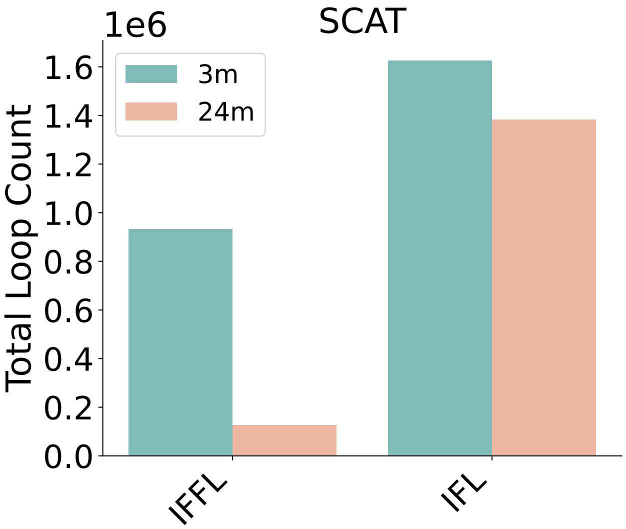


**Fig. S38. SCAT.**

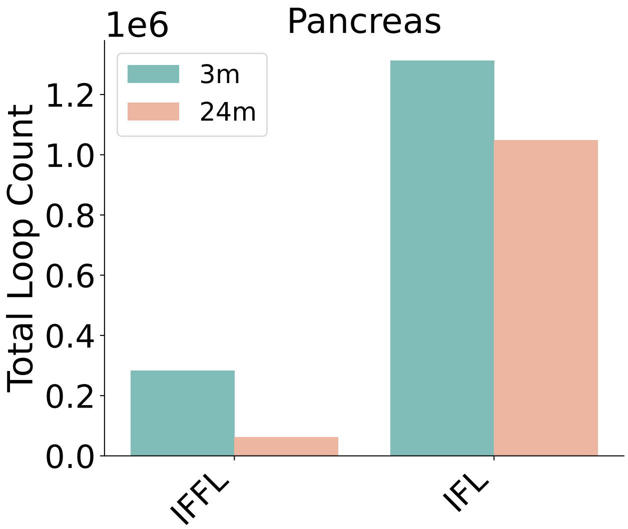


**Fig. S39. Pancreas.**

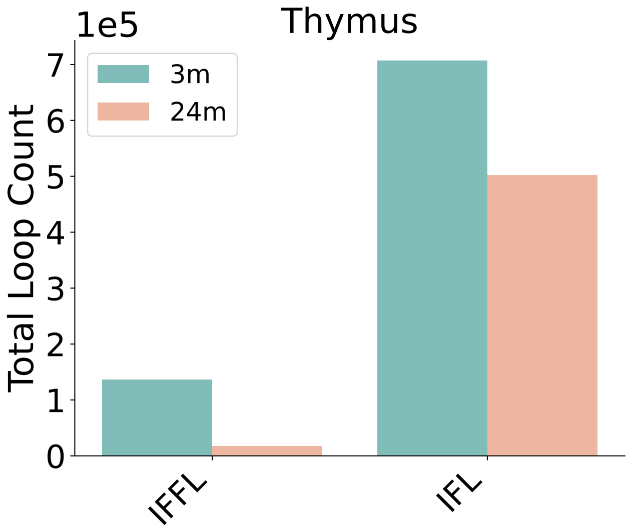


**Fig. S40. Thymus.**

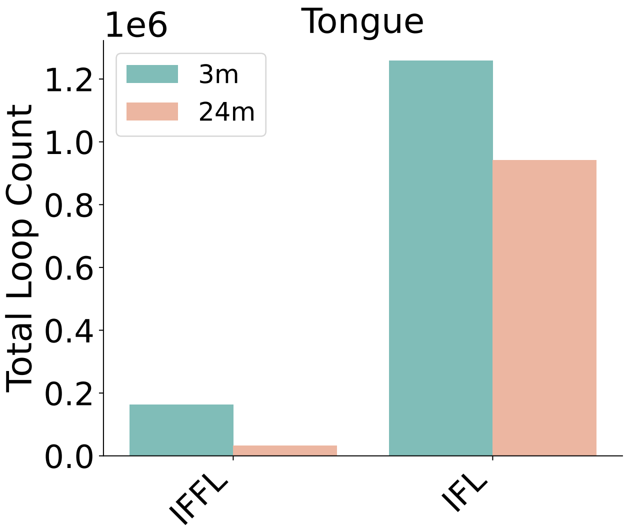


**Fig. S41. Tongue.**

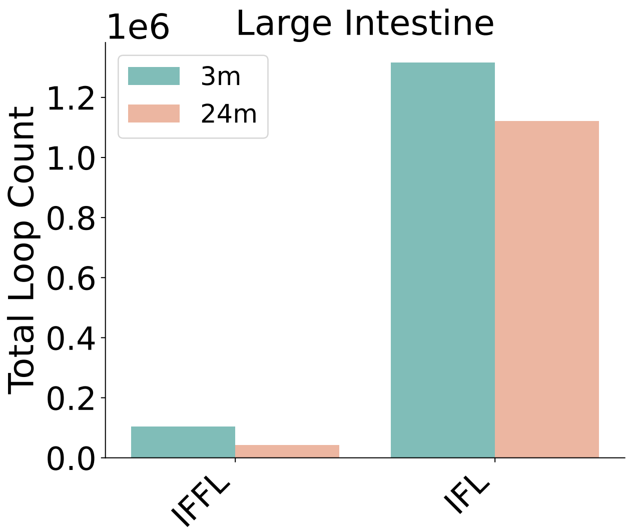


**Fig. S42. Large Intestine.**

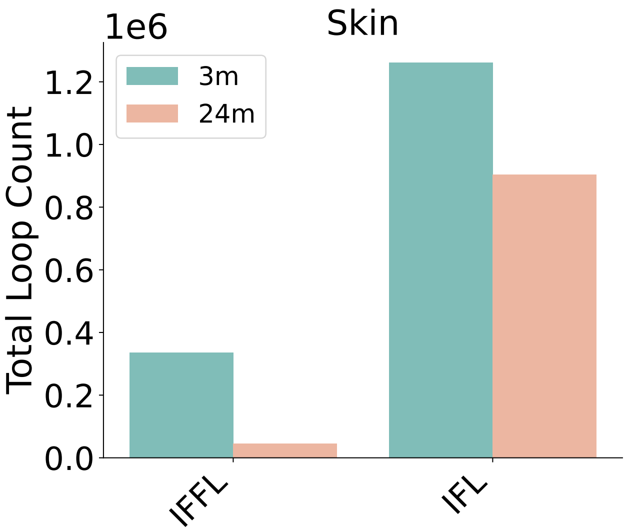


**Fig. S43. Skin 24m.**

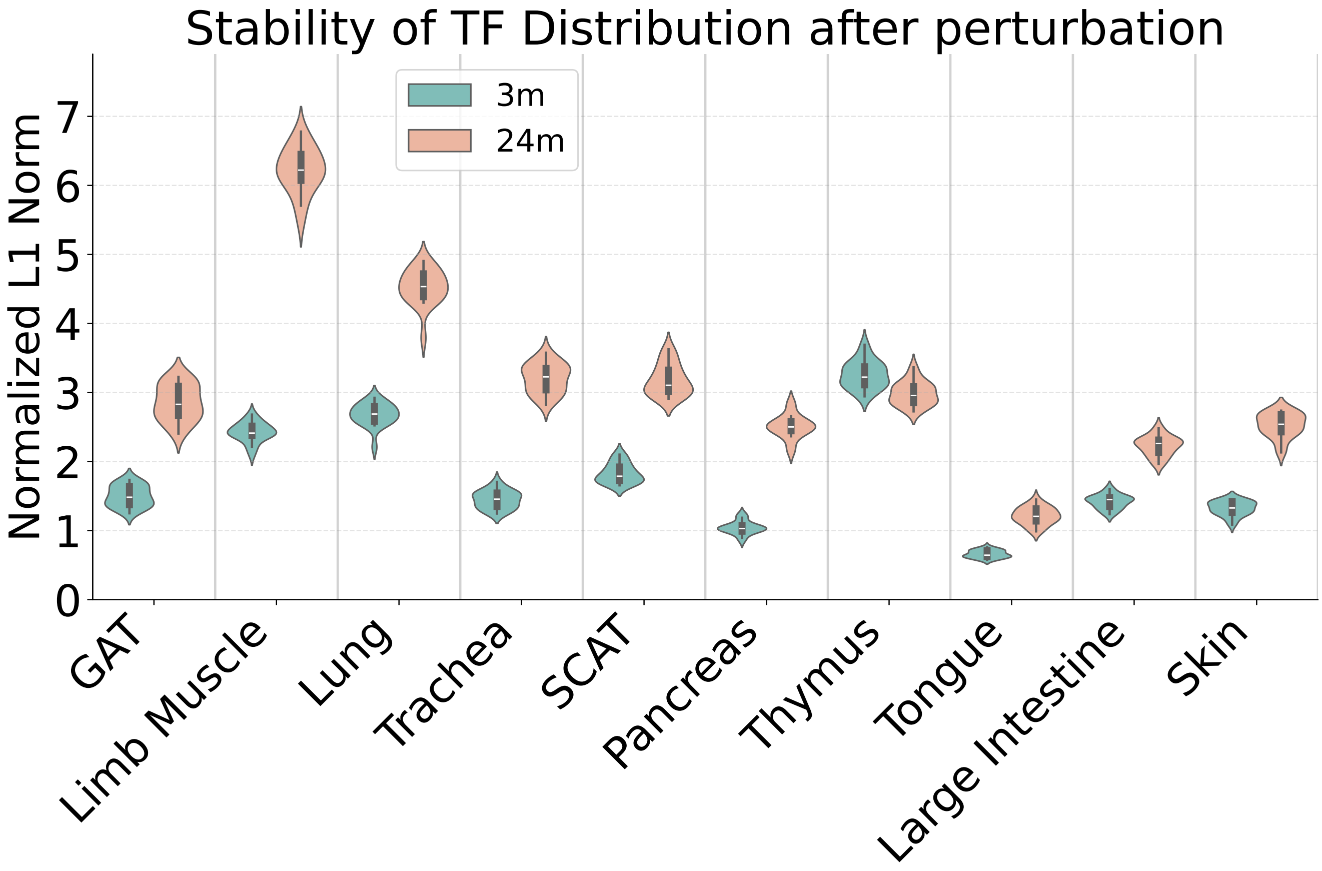


**Fig. S44.** The L1 Norm from perturbed TF Distribution to the original distribution, normalized by the mean TF activity in the data
